# PP4-dependent HDAC3 dephosphorylation discriminates between axonal regeneration and regenerative failure

**DOI:** 10.1101/446963

**Authors:** Arnau Hervera, Luming Zhou, Ilaria Palmisano, Eilidh McLachlan, Guiping Kong, Thomas Hutson, Matt C Danzi, Vance P. Lemmon, John L. Bixby, Andreu Matamoros-Angles, Kirsi Forsberg, Francesco De Virgiliis, Dina P. Matheos, Janine Kwapis, Marcelo A. Wood, Radhika Puttagunta, José Antonio del RÍo, Simone Di Giovanni

## Abstract

The molecular mechanisms discriminating between regenerative failure and success remain elusive. While a regeneration-competent peripheral nerve injury mounts a regenerative gene expression response in bipolar dorsal root ganglia (DRG) sensory neurons, a regeneration-incompetent central spinal cord injury does not. This dichotomic response offers a unique opportunity to investigate the fundamental biological mechanisms underpinning regenerative ability. Following a pharmacological screen with small molecule inhibitors targeting key epigenetic enzymes in DRG neurons we identified HDAC3 signalling as a novel candidate brake to axonal regenerative growth. *In vivo*, we determined that only a regenerative peripheral but not a central spinal injury induces an increase in calcium, which activates protein phosphatase 4 that in turn dephosphorylates HDAC3 thus impairing its activity and enhancing histone acetylation. Bioinformatics analysis of *ex vivo* H3K9ac ChIPseq and RNAseq from DRG followed by promoter acetylation and protein expression studies implicated HDAC3 in the regulation of multiple regenerative pathways. Finally, genetic or pharmacological HDAC3 inhibition overcame regenerative failure of sensory axons following spinal cord injury. Together, these data indicate that PP4-dependent HDAC3 dephosphorylation discriminates between axonal regeneration and regenerative failure.

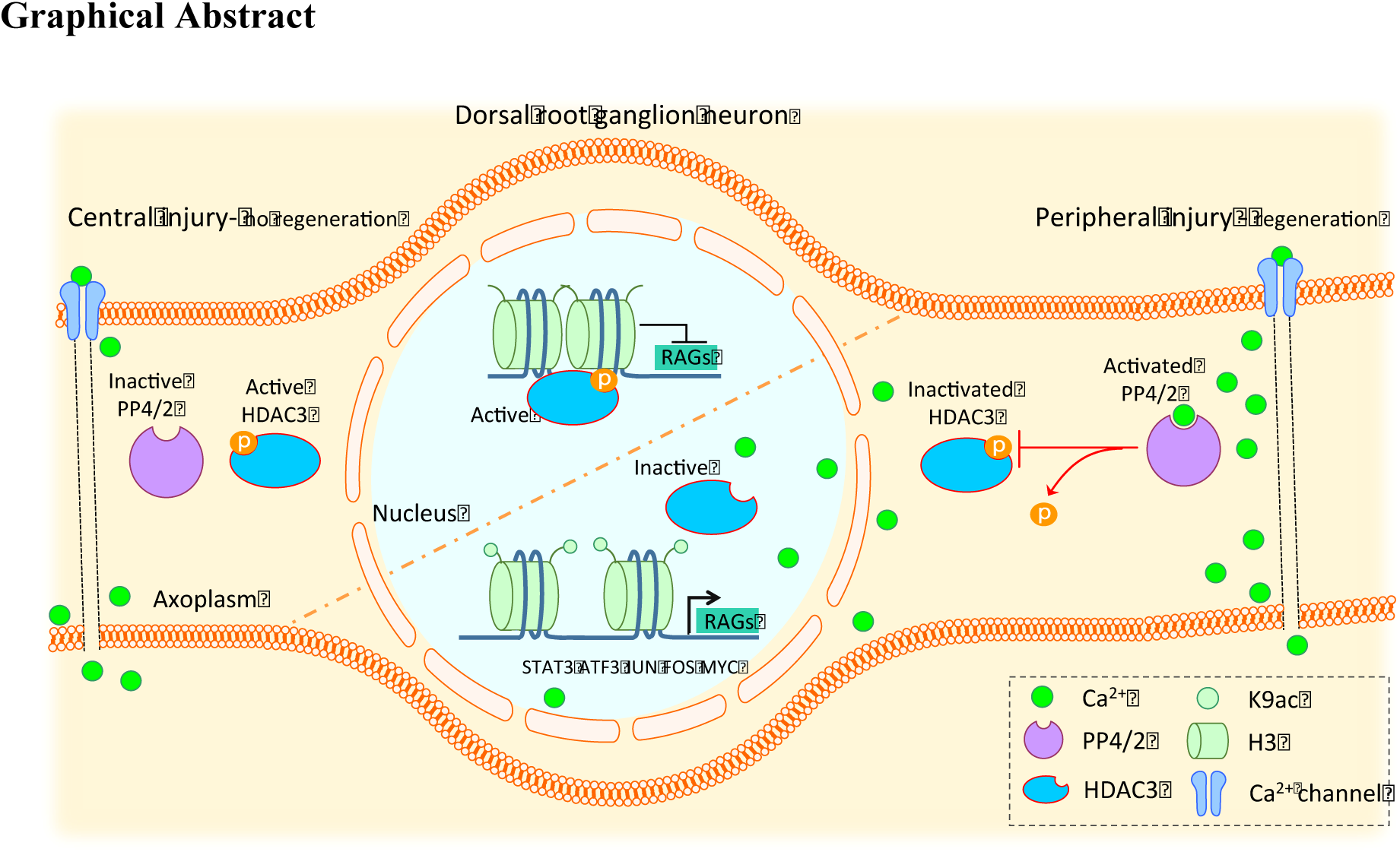
Graphical Abstract. Following central nervous system (CNS) spinal injury, protein phosphatase 4/2 activity is not induced since calcium levels remain unchanged compared to uninjured conditions. HDAC3 remains phosphorylated and occupies deacetylated chromatin contributing to its compaction inhibiting gene expression. Following peripheral nervous system (PNS) sciatic injury, protein phosphatase 4/2 activity is induced by calcium. HDAC3 is dephosphorylated leading to its inhibition and release from chromatin sites contributing to increase in histone acetylation and in the expression of regeneration associated genes (RAGs).

## Introduction

Following a central nervous system (CNS) injury, such as stroke or spinal cord injury, axonal regeneration is highly restricted. In stark contrast, spontaneous albeit partial functional axonal regeneration is possible after a peripheral nervous system (PNS) injury. This is likely due to the differences of the CNS and PNS in both intrinsic properties of neurons and the surrounding cellular environment. Importantly, glial cell-dependent signalling can affect the intrinsic properties of neurons (*1*), as is evident by the fact that the neuronal regenerative gene expression programme is only activated when axons lie within the PNS, but not in the CNS. This is typically modeled by the lumbar dorsal root ganglion (DRG) neurons (*2*, *3*), which project a peripheral axonal branch into the sciatic nerve (consisting of a permissive cellular environment) and a central axonal branch into the spinal cord (consisting of an inhibitory cellular environment). Strikingly, the peripheral branch of DRG mounts a robust regenerative response following a sciatic nerve injury, while the central branch fails to regenerate following a spinal injury (*2*, *4*). Furthermore, regeneration of the CNS branch is greatly enhanced by prior injury to the peripheral branch (conditioning lesion) (*2*, *4*) that leads to an increase in gene expression of a number of axonal growth and regeneration-associated genes, which does not occur after spinal lesions alone (*5*-*10*).

Dynamic gene expression changes are controlled and coordinated by epigenetic regulation such as DNA methylation or histone post-translational modifications which are essential in stem cell reprogramming, development of tissues and organs, as well as in cancer initiation (*11*). Strategies that take advantage of broad histone deacetlyase (HDAC) inhibitory drugs (class I and II such as TSA or MS-275) have been poor in promoting axonal regeneration after an optic nerve crush (*12*) or in enhancing regeneration past the lesion site after a spinal cord injury(*13*). Recently, we found that ERK-dependent phosphorylation leads to acetylation of histone H3K9 at the promoters of select RAGs after sciatic injury (*14*) and that viral overexpression of P300/CBP associated factor (P/CAF) promotes axonal regeneration after spinal cord injury by reactivating a regenerative gene expression programme (*14*). We also recently investigated DNA methylation in DRG following sciatic nerve versus spinal injury by methylation arrays including the CpG island rich promoter region of 13,000 genes and found that DNA methylation was not differentially represented on regeneration associated genes (*15*). However, this study fell short of providing genome wide DNA methylation profiles, therefore it did not allow ruling out a role for DNA methylation in axonal regeneration.

However, whether specific axonal signalling mechanisms can discriminate between axonal regeneration and regenerative failure by differentially activating or restricting the regenerative programme remains elusive. Here we hypothesize that critical axonal signals are conveyed to modify gene expression to restrict the regeneration programme after a non-regenerative spinal lesion, while these restrictions are lifted following a regenerative peripheral injury. Since regulation of gene expression relies on enzymes whose function is controlled by their activity, initially we adopted a pharmacological small molecule inhibitor screen of key epigenetic enzymes aiming to identify signalling pathways that might influence the regenerative growth potential of primary adult neurons and overcome regenerative failure. From this screen, we discovered that HDAC3 inhibition promoted neurite outgrowth on both growth permissive and inhibitory substrates. Next, we found that a regeneration-competent sciatic nerve injury induces an increase in calcium that activates protein phosphatase 4 (PP4) and slightly increases the activity of protein phosphatase 2 (PP2), which in turn dephosphorylate HDAC3, inhibiting its activity. However, a spinal lesion does not elicit increases in calcium signalling nor PP activity therefore failing to dephosphorylate HDAC3. Inhibition of PP4/2 activity promotes DRG regenerative growth similarly to mimicking HDAC3 phosphorylation.

Combined bioinformatics analysis of H3K9ac ChIPseq and RNAseq from DRG as well as HDAC3 protein-protein interaction databases suggested that HDAC3 might restrict the regenerative programme. Indeed, we found that inhibition of HDAC3 activity engages several regeneration-associated signalling pathways. Finally, we translated these findings to *in vivo* models of spinal cord injury where we found that either AAV-mediated overexpression of HDAC3 deacetylase-dead mutant or pharmacological HDAC3 inhibition overcome regenerative failure by promoting axonal growth of sensory axons. In summary, we found that calcium activation of PP4 leading to HDAC3 dephosphorylation discriminates between axonal regeneration and regenerative failure.

## Results

### Pharmacological inhibition and genetic manipulation show that HDAC3 inhibition selectively enhances neurite outgrowth on both growth permissive and non-permissive substrates

We performed a compound screen to identify whether inhibiting the activities of enzymes that modify key epigenetic marks could promote neurite outgrowth of adult neurons on both growth permissive and non-permissive inhibitory substrates. All inhibitors were initially screened at various concentrations in a neurite outgrowth assay using adult mouse dorsal root ganglia (DRG) neurons cultured on a growth permissive laminin-PDL substrate. The inhibitors that promoted significant outgrowth were further examined on a myelin growth inhibitory substrate.

We screened 14 pharmacological inhibitors of enzymes affecting epigenetic marks (Table 1). These include inhibitors of DNMT1-3, DNMT 3b, HDAC class II (HDAC 4, 5, 7, 9), HDAC 1, HDAC 1-2, HDAC3, the HMT G9a for H3K9me3, EZH2 for H3K27me3, DOT1L for H3K79me3, the HKDM JMJD2 for H3K9me3, JMJD3 for H3K27me3, and the HDR PADI4 for H3R2-8-17cit. Inhibitors of bromodomain BRD2/3/4 and BAZ2 proteins that are linked to both permissive and repressive epigenetic marks were also examined. The EZH2 inhibitor *43A* and the HDAC3 inhibitor *RGFP966* significantly enhanced neurite outgrowth by more than 2 fold compared to vehicle. However, only *RGFP966-dependent* HDAC3 inhibition resulted in significant and extensive neurite outgrowth on both permissive laminin and inhibitory myelin substrates (Figure 1A-C), and enhanced H3K9 acetylation as expected (Figure 1D, E). In contrast, inhibitors of HDAC class II or HDAC 1 (233) and HDAC1-2 (963) did not affect DRG outgrowth, although they increased overall histone acetylation as expected (Figure 1A, Supplementary Figure 1A, B). Next, we infected cultured DRG neurons plated on PDL/laminin or myelin with an AAV-HDAC3 deacetylase dead mutant (HDAC3mut), a control AAV-GFP or AAV-HDAC3wt. We found that while overexpression of wt HDAC3 restricted outgrowth, overexpression of mutant HDAC3 strongly promoted neurite outgrowth on both permissive and inhibitory substrates and enhanced histone acetylation (Figure 1F, G, Supplementary Figure 1C, D). Similarly, HDAC3 gene silencing with validated shRNAs promoted DRG outgrowth (Supplementary Figure 1 E, F), supporting the HDAC3 pharmacological inhibition and deacetylase mutant overexpression data. This indicates that HDAC3 may be a repressor of axonal regeneration whose inhibition may allow gene expression-dependent regenerative reprogramming.

**Table 1.** List of inhibitory drugs, target molecule and epigenetic mark employed in a screening for DRG neurite outgrowth.

| **Drug** | **Enzyme** | **Target** | **Function** |
| --- | --- | --- | --- |
| 5-AZA | DNMT1/3 | DNMT | 5'-methylcytosine |
| 13X | DNMT3B | DNMT | 5'-methylcytosine |
| 69A | HDAC class II (4/5/7/9) | HDAC | H3/4ac |
| 233 | HDAC1 | HDAC | H3/4ac |
| 963 | HDAC1/2 | HDAC | H3/4ac |
| 966 | HDAC3 | HDAC | H3/4ac |
| 801 | BAZ2A/ BAZ2B | HME/DNMT | H3/4ac |
| 51A | BRD 2/3/4 | S/T kin | H3/4ac |
| 17A | DOT1L | HMT | H3K79me3 |
| J4/77A | JMJD3 | HKDM | H3K27me3 |
| 71B | G9A | HMT | H3K9me3 |
| 484 | PADI4 | HDI/HRDM | H3R(2/8/17)cit |
| 67A | JMJD2/JARID1 | HKDM | H3K9me3/36me3/4me3 |
| 43A | EZH2 | HMT | H3K27me3 |

**Figure 1.**
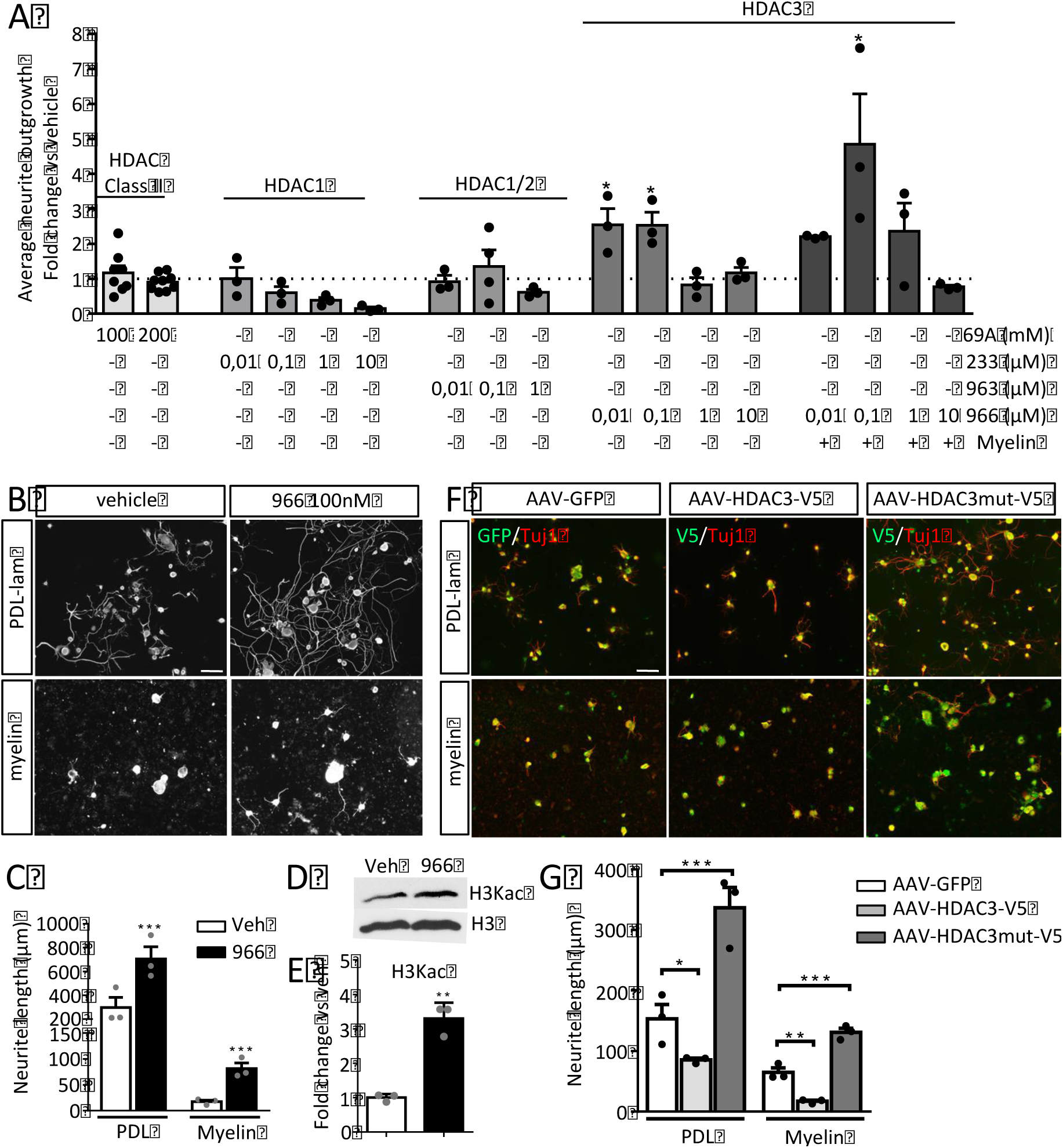
Pharmacological HDAC3 inhibition and HDAC3 dead mutant enhance DRG neurite outgrowth. A. RGFP966 (966)-dependent HDAC3 inhibition resulted in significant and robust neurite outgrowth compared to HDAC class II, HDAC1/2 and HDAC1 or 3 inhibition via 69A, 963 and 233 respectively. B, C. Specific pharmacological inhibition of HDAC3 facilitates DRG neurite outgrowth (average neurite outgrowth, Tuj1 positive neurons) on PDL-laminin or Myelin substrates. D, E. Immunoblotting shows significantly increased H3K9 acetylation after 966 vs vehicle. Scale bars, 100 μm. F, G. AAV-mediated mutant HDAC3 (Y298H, V5) infection of DRG neurons induces neurite outgrowth (Tuj-1/V5 double positive neurons), which is repressed by wildtype HDAC3 overexpression. Scale bars, 50 μm. For all graphs, data expressed as mean fold change or average neurite length ± s.e.m. N= 3 or 5 biological replicates. (*p<0.05, **p<0.01, ***p<0.005) indicate significant difference of 966 versus vehicle or V5 vs AAV-GFP (C and E, Student’s t-test) (A and G, ANOVA followed by Bonferroni test).

### HDAC3 phosphorylation and activity are inhibited by a peripheral sciatic but not by a central spinal injury

Next, we investigated the post-injury regulation of HDAC3 in the pseudo-unipolar DRG system *in vivo* following a regeneration-competent peripheral sciatic nerve axotomy (SNA) injury versus a regeneration-incompetent central spinal axotomy (DCA). This allowed for the comparison of HDAC3 regulation in the same neuronal population undergoing a regenerative versus non-regenerative axonal lesion. HDAC3 can be regulated at the gene or protein expression level; it can be shuttled between cytoplasm and nucleus; and it can be phosphorylated (*16*-*18*). First, we measured the gene expression of HDAC3 and of the other class I HDACs, HDAC1 and 2 by qPCR, in sham versus axonal injury in DRG. We found that all HDACs are expressed in DRGs, but neither a peripheral nor a central injury modifies their gene expression level (Supplementary Figure 2 A, B). Protein analysis of HDAC3 expression revealed that it is highly expressed in DRG neurons, mainly in the nucleus, but again neither its protein expression nor its nuclear-cytoplasmic shuttling was regulated by either a sciatic or spinal injury (Supplementary Figure 3A-E). However, we found that the nuclear phosphorylation of HDAC3 (Ser424) was strongly reduced by a sciatic but not by a central spinal injury (Figure 2A-E). Since phosphorylation of HDAC3 promotes its enzymatic activity, we measured the activity of HDAC3 in DRG after DCA or SNA to find that HDAC3 deacetylase activity was strongly reduced by SNA only (Figure 2F), in line with its reduced phosphorylation.

**Figure 2.**
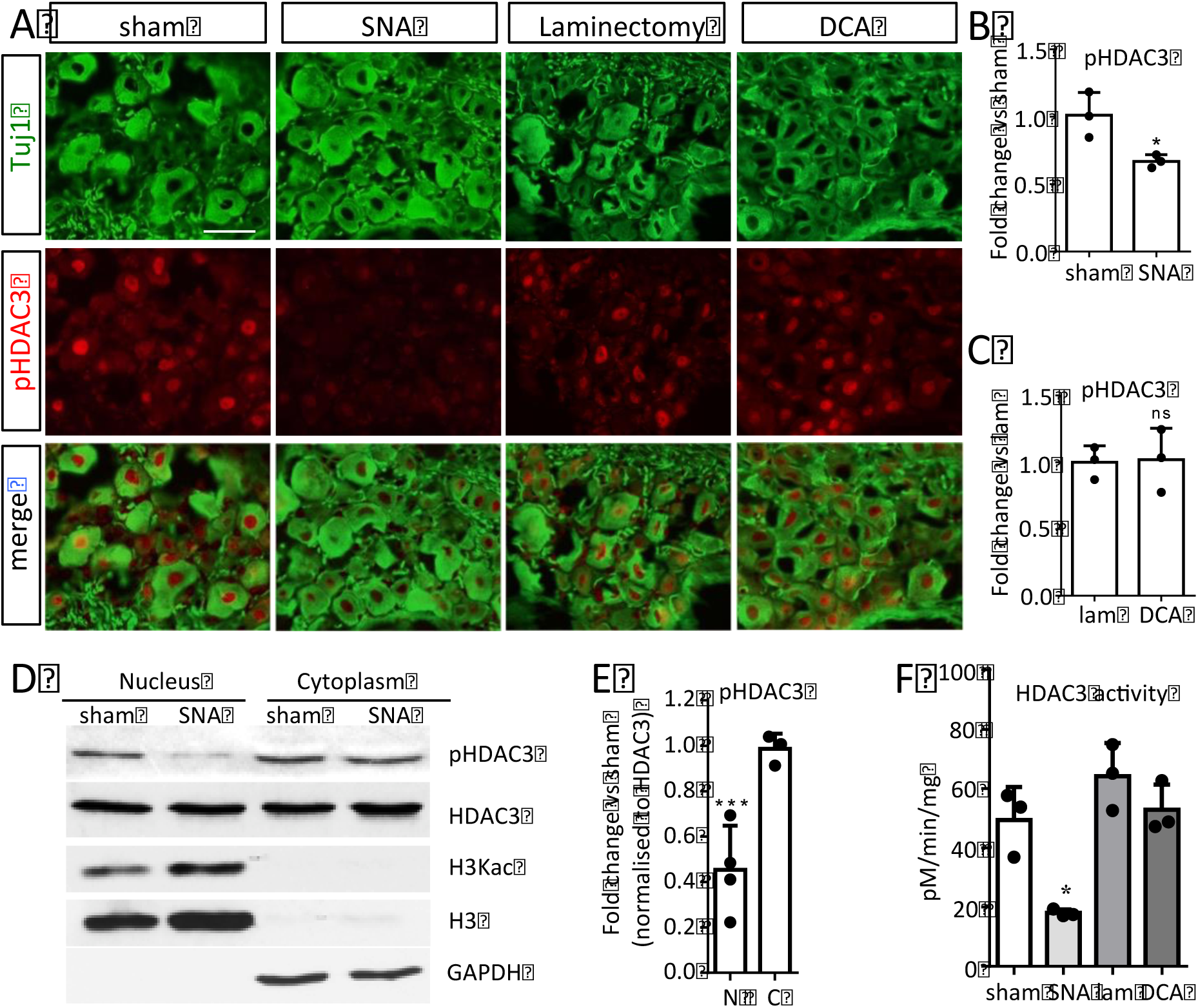
Regenerative sciatic nerve injury selectively decreases HDAC3 phosphorylation and activity in DRG neurons. A. IHC in DRG sections for pHDAC3 revealed that HDAC3 phosphorylation is decreased, primarily in the nucleus 24h after sciatic nerve axotomy (SNA) but not spinal dorsal column axotomy (DCA). Scale bars, 100 μm. B-C. Bar graphs show mean fold change ± s.e.m. N= 3 animals. *p<0.05 indicate significant difference versus sham or laminectomy (lam) (Student’s t-test). D. Western blot revealed that HDAC3 phosphorylation is decreased in DRG nuclear extracts but not in the cytoplasm 24h after SNA.E. Bar graphs show intensity measurement of nuclear vs cytoplasm immunoblot. Data is expressed as mean ± s.e.m. N= 3. (***p<0.005) indicate significant difference versus sham (Student’s t-test). F. HDAC3 activity (HDAC3 IP and activity assay) is strongly reduced after SNA but not DCA. Data is expressed as mean ± s.e.m. N= 3 animals. (*p<0.05) indicate significant difference versus sham (ANOVA followed by Bonferroni test).

**Figure 3.**
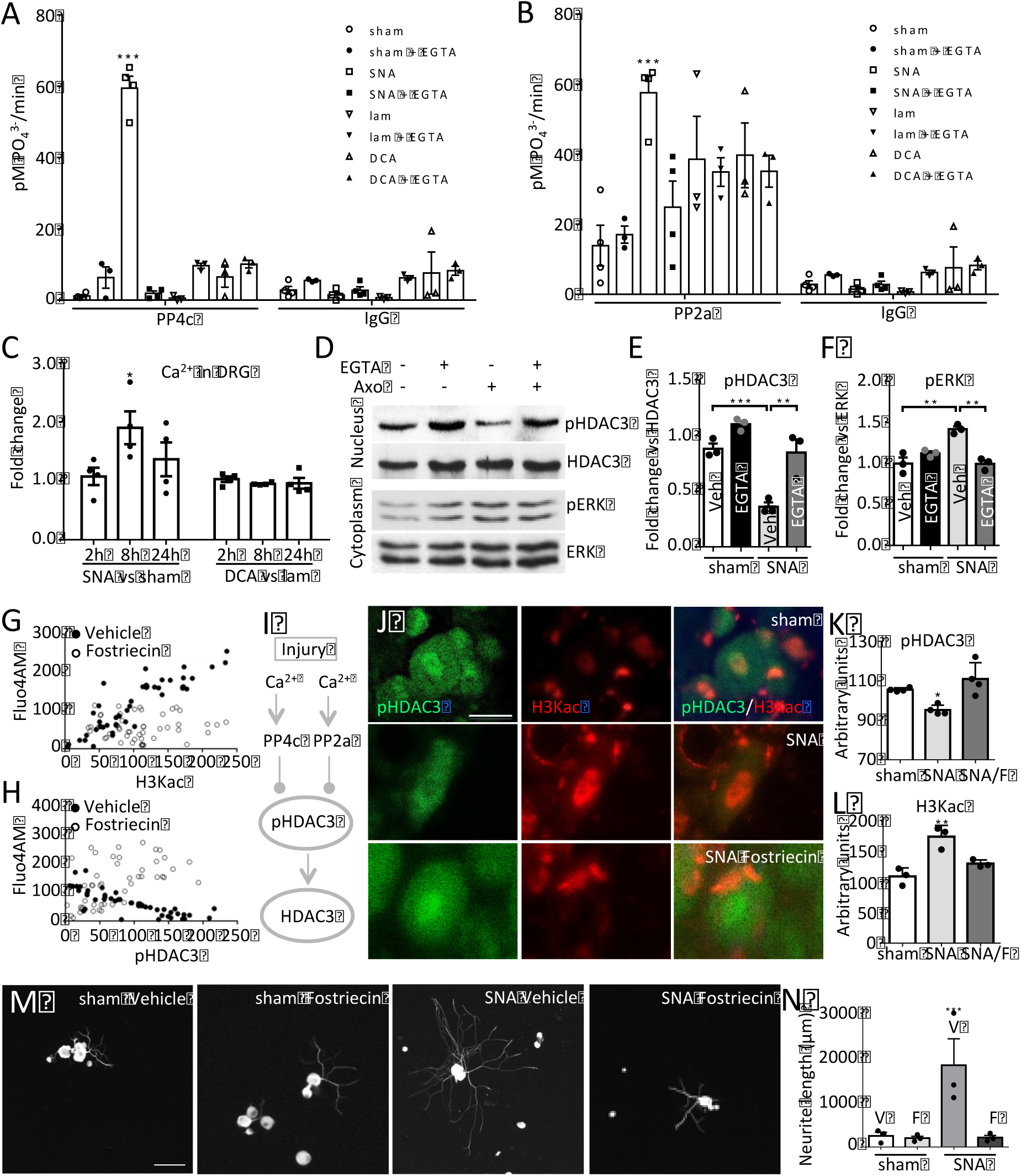
SNA induced calcium regulates PP4/2 activity that in turn control HDAC3 phosphorylation and DRG regenerative growth. A-B. PP4 and PP2 activities are induced after SNA but not DCA, and are blocked by the calcium chelator EGTA. Data is expressed as mean ± s.e.m. N= 3-4 animals. (***p<0.001) indicate significant difference versus sham or lam (Two-way ANOVA followed by Bonferroni test). C. SNA but not DCA significantly enhances calcium levels in DRG. Data is expressed as mean ± s.e.m. N= 4 animals. (*p<0.05) indicate significant difference versus sham or lam (ANOVA followed by Bonferroni test). DF. Immunoblotting of DRG homogenates shows that nerve injury-dependent nuclear HDAC3 dephosphorylation is reduced by the calcium scavenger EGTA upon *in vivo* delivery on the injured sciatic nerve (Axo=Axotomy). Cytoplasmic pERK levels are used as control of axotomy-dependent signalling. E-F. Data is expressed as mean fold change of band intensity levels ± s.e.m. N= 3. (**p<0.01, ***p<0.001) indicate a significant difference (ANOVA followed by Bonferroni test). G-H. Significant Pearson correlations between calcium levels (Fluo4AM) and H3K9ac, r^2^: 0.7802 (G); or pHDAC3, r^2^: 0.6366 (H) in cultured DRG neurons after KCl. The Pearson correlations were disrupted by administration of Fostriecin (calcium and H3K9ac, r^2^: 0.0007; calcium and pHDAC3, r^2^: 0.3025). Data is expressed as single cell fluorescence levels. N=50 cells per condition from 4 biological replicates. I. Schematic pathway summarizing Ca^2+^ and PP4c and PP2a-dependent HDAC3 dephosphorylation. J-L. Immunofluorescence (J) in DRG tissue sections 24h after injury *in vivo* shows that the PP2a/PP4 inhibitor Fostriecin (F) inhibits the injury-induced decrease in pHDAC3 (K) and the increase in H3K9ac (L). Scale bar, 10 μm. K-L. Data is expressed as mean fluorescence intensity in arbitrary units ± s.e.m. N= 3 animals (*p<0.05; **p<0.01) indicate significant difference versus sham (ANOVA followed by Bonferroni test). M-N. Pharmacological inhibition of PP2a/PP4 impairs conditioning-induced DRG neurite outgrowth *ex vivo.* M. DRG neurite outgrowth (TUJ1 positive cells) 24h after sham or sciatic nerve axotomy (SNA) prior to i.t. administration of vehicle (V) or Fostriecin (F) (240μM). Scale bar, 100μm. N. Data is expressed as average neurite length per neuron ± s.e.m. N= 3 animals per condition, technical triplicate. *** p<0.005 indicate significant difference versus sham/vehicle (One way ANOVA followed by Bonferroni).

### Sciatic nerve but not spinal injury induces calcium-dependent PP4 activity that is required for HDAC3 dephosphorylation

Calcium is a prominent second messenger triggered by the regenerative sciatic nerve injury, although whether it fails to be induced in DRG by a central spinal injury remains elusive. Protein phosphatases 4c and 2a (PP4c and PP2a), are activated via calcium signalling, and have been shown to interact with HDAC3. However only PP4c has been shown to dephosphorylate HDAC3 in cancer cells (*16*, *18*), nonetheless whether PP4c and PP2a control HDAC3 phosphorylation in neurons has not been investigated. Therefore, we hypothesised that a regenerative peripheral SNA but not a central DCA would lead to calcium-dependent activation of PP4c and/or PP2a. This would in turn inhibit HDAC3 via dephosphorylation resulting in an increase in histone acetylation in DRG neurons. First, we measured the activity of PP4c or PP2a following SNA or DCA. We found that SNA but not DCA strongly promoted calcium-dependent PP4c and PP2a activity that was inhibited by administration of the calcium chelator EGTA (Figure 3A, B). Accordingly, we also found that calcium, required for PP activity, was induced by SNA but remained unchanged following DCA (Figure 3C). While PP4c basal activity was very low in DRG and was strongly induced by SNA (Figure 3A), PP2a basal levels were higher (Figure 3B), suggesting that PP4c is the preferentially inducible protein phosphatase selectively following SNA. Moreover, the calcium chelator EGTA was able to block the injury induced increase in PP4c and PP2a activity, highlighting the calcium dependence of the phosphatase activity induced by SNA. We next investigated whether phosphorylation of HDAC3 in DRG was also dependent upon calcium. Indeed, we found that HDAC3 dephosphorylation after sciatic nerve injury required calcium, as administration of EGTA on the nerve at the time of injury blocked nuclear HDAC3 dephosphorylation without altering HDAC3 expression (Figure 3D-F). In support of these findings, we observed a significant correlation between HDAC3 dephosphorylation but not total HADC3 and calcium signal intensity after potassium chloride (KCl)-dependent calcium induction in DRG neurons (Supplementary Figure 4 A-C and D-E). Additionally, when we inhibited PP4/2a activity with the PP4/2a inhibitor Fostriecin in KCl treated DRG cells, we observed that the correlation between calcium and HDAC3 dephosphorylation or histone acetylation was lost (Figure 3G, H). Next, we asked whether PP4/2a activity was required for HDAC3 dephosphorylation and for DRG regenerative growth. We delivered PP4/2a inhibitor Fostriecin or vehicle intrathecally immediately after sham or SNA, 24 hours later we measured HDAC3 phosphorylation and H3K9ac in DRG or performed a neurite outgrowth assays in cultured DRG neurons. Importantly, first we found that PP4/2a inhibition led to an increase in HDAC3 phosphorylation with a concomitant decrease in H3K9ac in DRG neurons (Figure 3I-L). Next, we observed that PP4/2a inhibition blocked SNA-dependent DRG regenerative growth (Figure 3M, N). When we silenced PP4c or PP2a by electroporation of siRNA against PP4c or PP2a in KCl treated DRG cells, we observed that KCl-dependent HDAC3 dephosphorylation was significantly reduced (Figure 4A-G). Antibodies to PP2a were available for western blotting and antibodies to PP4 were available for immunocytochemistry only, determining the choice of the protein detection system including after silencing. We finally asked whether HDAC3 serine 424 phosphorylation (ser424) that controls HDAC3 activity and histone acetylation (*19*), is able to regulate DRG outgrowth. We overexpressed phospho-mimetic or phospho-dead HDAC3 S424A or S424D mutant plasmids in cultured DRG cells respectively. As expected, while HDAC3 phospho-dead S424A significantly promoted DRG outgrowth compared to control vector or HDAC3 WT, phosphor-mimetic HDAC3 S424D inhibited it (Figure 4H, I). All together, these data suggest that peripheral nerve injury induces calcium that is required for PP4c and to a lesser extent PP2a activity. PP4/2a are in turn needed to inhibit HDAC3 activity through dephosphorylation and to increase histone acetylation leading to DRG regenerative growth, including after a conditioning lesion. Lastly, HDAC3 serine 424 dephosphorylation, which requires PP4/2a, specifically controls DRG regenerative growth.

**Figure 4.**
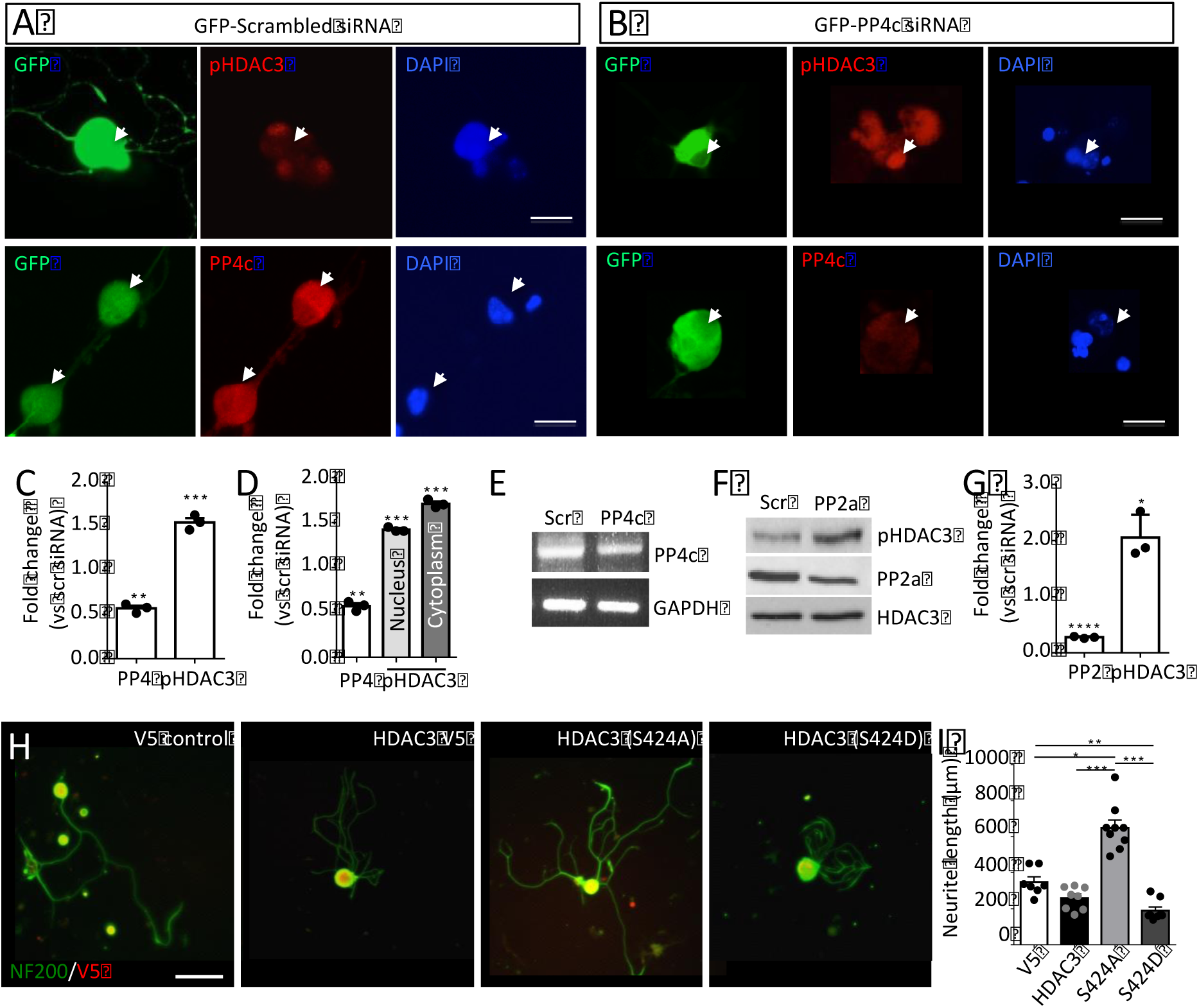
PP4c and PP2A dephosphorylate HDAC3 at Ser424, which is required for
DRG outgrowth. A-E. PP4c gene silencing promotes HDAC3 phosphorylation in the presence of KCl (A-C, arrows), both in the nucleus and the cytoplasm (D). Scale bar, 10 μm.C-D. Data is expressed as mean fold change vs scrambled (scr) siRNA of fluorescent intensity of pHDAC3 or PP4c in GFP positive cells ± s.e.m. N=3. (**p<0.01; ***p<0.005) indicate significant difference versus scrambled siRNA (Student’s t-test). E. PCR against PP4c demonstrates silencing of PP4c mRNA. F-G. PP2a gene silencing inhibits the KCl induced dephosphorylation of HDAC3 as shown by immunoblotting of pHDAC3 (F). G. Data is expressed as mean fold change of immunoblot band intensity ± s.e.m. N=3 (*p<0.05; ***p<0.005) indicate significant difference versus control siRNA (Student’s t-test). H-I. Genetic modification of activity-dependent phosphorylation of serine 424 of HDAC3 affects DRG neurite outgrowth. H. DRG neurite outgrowth after transfection of control (V5 control), WT HDAC3-V5, phospho-dead HDAC3 (S424A)-V5 or phospho-mimetic HDAC3 (S424D)-V5. Scale bar, 50μm. I. Data is expressed as mean ± s.e.m. N= 3, approximately 35 V5 positive cells each. * p<0.05, **p<0.01, *** p<0.005 indicate significant difference (One way ANOVA followed by Bonferroni).

**Figure 5.**
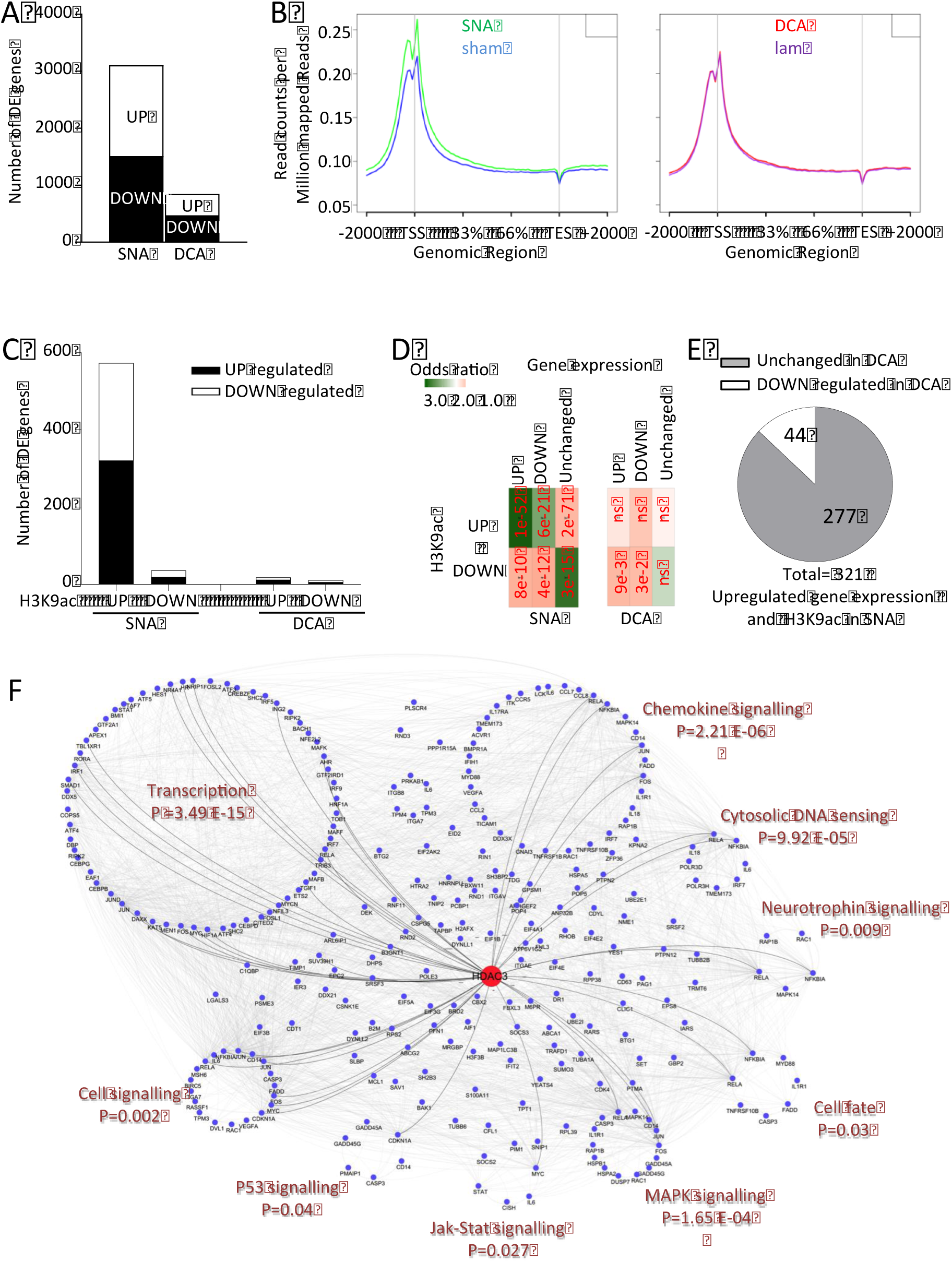
H3K9ac ChIPseq and HDAC3 interactome signalling network. A. The graph shows the number of differentially expressed (DE) genes 24h following SNA or DCA compared to their respective controls. B. RNAseq read counts show differential occupancy of H3K9ac on specific gene regions, including an increased occupancy close to the transcriptional start site (TSS) after SNA only, when compared to corresponding controls. C. Bar graph shows the number of genes that are upregulated or downregulated following SNA or DCA (RNAseq) with an increased or decreased H3K9ac occupancy (ChIPseq), respectively. D. Odds ratio analysis of the enrichment/depletion of genes with differential H3K9ac occupancy across each upregulated and downregulated gene cluster in (C); the numbers in red represent the *p* value given by Fisher’s exact test (ns= not significant). E. Out of the 321 upregulated genes associated with increased H3K9ac upon SNA, the majority does not show a change in expression following DCA (grey), while some of them are downregulated (white). F. Cytoscape visualization of the protein network resulting from HDAC3 interactome and H3K9ac ChIPseq and RNAseq upon SNA. The signalling network was created by combining the protein interactome of HDAC3 and H3K9ac dependent genes with increased gene expression upon SNA.

### Combinatorial bioinformatics analysis of RNAseq, H3K9ac ChIPseq and HDAC3 protein-protein interactome suggest that HDAC3 is involved in the control of the regenerative gene expression programme

So far, we found that pharmacological or genetic HDAC3 inhibition (HDAC3mut) enhanced DRG neurite growth as well as histone H3K9 acetylation and that regeneration-competent SNA induces HDAC3 dephosphorylation and histone H3K9 acetylation in DRG neurons. Since H3K9ac depends on HDAC3 activity and is enriched at active promoters controlling gene expression, we investigated whether a regeneration-competent SNA would be associated with increased gene expression and H3K9ac enrichment. Therefore, we generated and combined RNAseq ((*20*)GSE97090, (https://www.ncbi.nlm.nih.gov/geo/query/acc.cgi?token=anmdoqiodzgfvat&acc=GSE97090) with ChIPseq for H3K9ac (GSE108806 (exwheyyitzqlpoz)) from sciatic DRGs 24 hours after regenerative SNA vs non-regenerative DCA. RNA sequencing identified 3090 differentially regulated genes after SNA (1503 upregulated, 1587 downregulated). Whilst after DCA, the gene expression programme was decreased by 3.7 fold (471 upregulated, 363 downregulated) (*p* value<0.05) (Figure 5A). H3K9ac ChIP revealed a significant enrichment of recovered DNA versus control IgG ChIP (Supplementary Figure 5A). As expected, enrichment of H3K9ac after ChIP sequencing was prevalent at TSS, enhancers and gene bodies (Supplementary Figure 5B). Genome wide analysis of H3K9ac revealed that H3K9ac occupancy increased specifically after SNA with respect to sham, but not after DCA with respect to laminectomy control (Figure 5B). This prompted us to assess the gene expression programme associated with changes in H3K9ac occupancy following SNA. To this end, we integrated ChIPseq and transcriptome data that allowed identifying 573 genes with increased H3K9ac occupancy after SNA (TSS+/-1000bp plus gene body) associated with changes in gene expression (321 upregulated and 252 downregulated, *p* value<0.05), while only few genes showed changes in H3K9ac occupancy following DCA (Figure 5C and Supplementary File 1). Increased H3K9ac occupancy was more highly correlated with upregulated genes (Fisher exact P=1e-52, Figure 5D), decreased occupancy correlated with non-differentially expressed genes (Fisher exact P=3e-15, Figure 5D), supporting the interplay between enhanced histone acetylation and gene expression. It is worth noting that upon DCA, differentially expressed genes are significantly depleted for transcripts with changes in H3K9ac occupancy (Figure 5D). We next assessed whether these 321 upregulated SNA-responsive genes associated with increased H3K9ac occupancy were affected by nonregenerative DCA. Interestingly, we found that all of these transcripts were either downregulated or that their expression did not change (Figure 5E and Supplementary File 1). The opposing activation of these H3K9ac-dependent genes between the two injury paradigms suggests that this might contribute to determining whether axonal regeneration is induced or not.

Next we asked whether i) the HDAC3 interactome (network of HDAC3 proteinprotein interactions) could be regulated after injury and whether ii) it shared key signalling pathways with those found to be associated with axonal regeneration. Therefore, we generated a HDAC3 interactome by FpClass analysis (Supplementary File 2) and we evaluated how many of these HDAC3 interactors were differentially regulated after injury. Interestingly, about 23% of HDAC3 interactors were differentially regulated after SNA with 12.7% being upregulated and 9.8% downregulated, while only about 4% were differentially regulated after DCA (Supplementary Figure 5C). GO analysis showed that the SNA-differentially regulated HDAC3 interactors are mainly involved in transcriptional regulation (Supplementary File 3). Indeed, noteworthy, many of those SNA-upregulated HDAC3 interactors (Supplementary Figure 5D, E, green) are regeneration-associated transcription factors such as Atf3, Jun, Fos, Cebpd, Klf6, Hif1a. Some of them showed opposite regulation after DCA (Supplementary Figure 5D). Moreover, KEGG pathway analysis specifically revealed that HDAC3 interactors that are upregulated following SNA are enriched for signalling pathways associated with axonal regeneration, while this is not the case for the ones that are downregulated (Supplementary Figure 6A, B and Supplementary File 3). These pathways include MAPK, JAK-STAT3, neurotrophin, PI3K/AKT, cAMP signalling.

**Figure 6.**
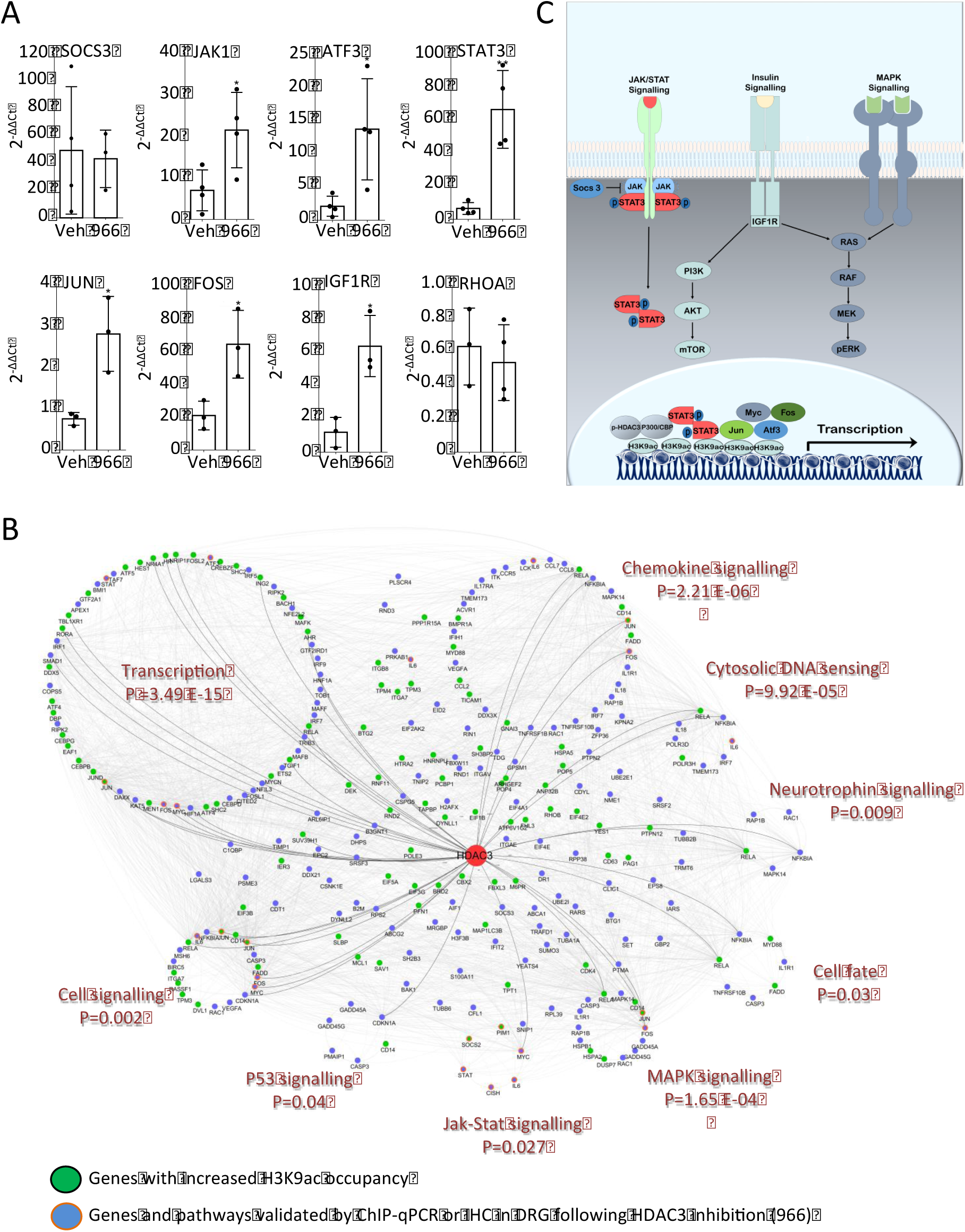
Intrathecal pharmacological inhibition of HDAC3 activity increases H3K9 acetylation on promoters of regenerative genes in DRG after DCA. A. ChIP-qPCR data following 966-dependent inhibition of HDAC3 after DCA (24 hours) shows an increase in H3K9ac on promoters of regeneration associated genes. Data is expressed as 2^−AACt^ normalized versus IgG and input for each sample ± s.e.m. N= 4 samples per condition (5 animals per sample). * p<0.05, ** p<0.01 indicate significant difference versus vehicle (Student’s t-test). B. Cytoscape visualization of the HDAC3-dependent network as shown in Figure 5. Here, highlighted in orange are the validated pathways and targets via ChIP-qPCR(Figure 6A) or IHC (Supp Figure 7). C. Schematic diagram showing HDAC3-dependent signalling pathways.

To identify possible upstream TFs that may be responsible for H3K9ac-dependent gene expression following SNA, we performed transcription factor enrichment analysis on the promoter of the identified 321 H3K9ac regulated genes (Supplementary File 4). Nineteen out of the identified 75 predicted TFs are found among the HDAC3 interactors which have been related to neuronal plasticity and regeneration (Supplementary Figure 6C, D).

To probe the potential transcriptional and downstream signalling events associated with HDAC3 reduced activity upon regenerative injury, we built a protein-protein interaction network of the genes that were upregulated after SNA with increased H3K9ac occupancy and generated a hypothetical HDAC3-responsive signalling network (Figure 5F). The network displayed significant enrichment for signalling involved in the axonal regeneration programme including transcription factors, JAK-STAT, p53, neurotrophin, Wnt, MAPK signalling pathways and actin cytoskeleton regulation (Figure 5F and Supplementary File 5).

Taken together, these data suggest that H3K9ac and HDAC3 may be involved in the control of the regenerative programme and that modulating HDAC3 activity might lead to synergistically targeting various signalling events that are associated with a regenerative phenotype. To verify this prediction, we investigated whether the *RGFP966*-dependent inhibition of HDAC3 following spinal injury (DCA) would enhance the expression level and promoter histone acetylation of key regenerative transcription factors and signalling pathways including the ones implied by our bioinformatics analysis (activated upon SNA but unchanged upon DCA) and summarized in the HDAC3-dependent signalling network in Figure 5F. We also tested where HDAC3 inhibition would be able to activate signalling pathways elicited by SNA and previously associated with axonal regeneration including JAK-STAT3, IGF1 and MYC. We delivered RGFP966 intrathecally at the time of DCA and 24 hours later we tested the expression and H3K9ac occupancy of transcription factors and signalling molecules in DRG. We found increased H3K9 acetylation on ATF3, JUN, FOS, JAK1, STAT3, IGF1R promoters in 966 vs vehicle treated DRG (Figure 6A). Consistently, we found an increased protein expression for ATF3, JUN, pSTAT3, MYC, IGF1R and pERK in injured DRG neurons, however not in non-neuronal DRG cells, 5 weeks following RGFP966 treatment and SCI (Supplementary Figure 7), at a time where we also measured regeneration of DRG sensory neurons.

**Figure 7.**
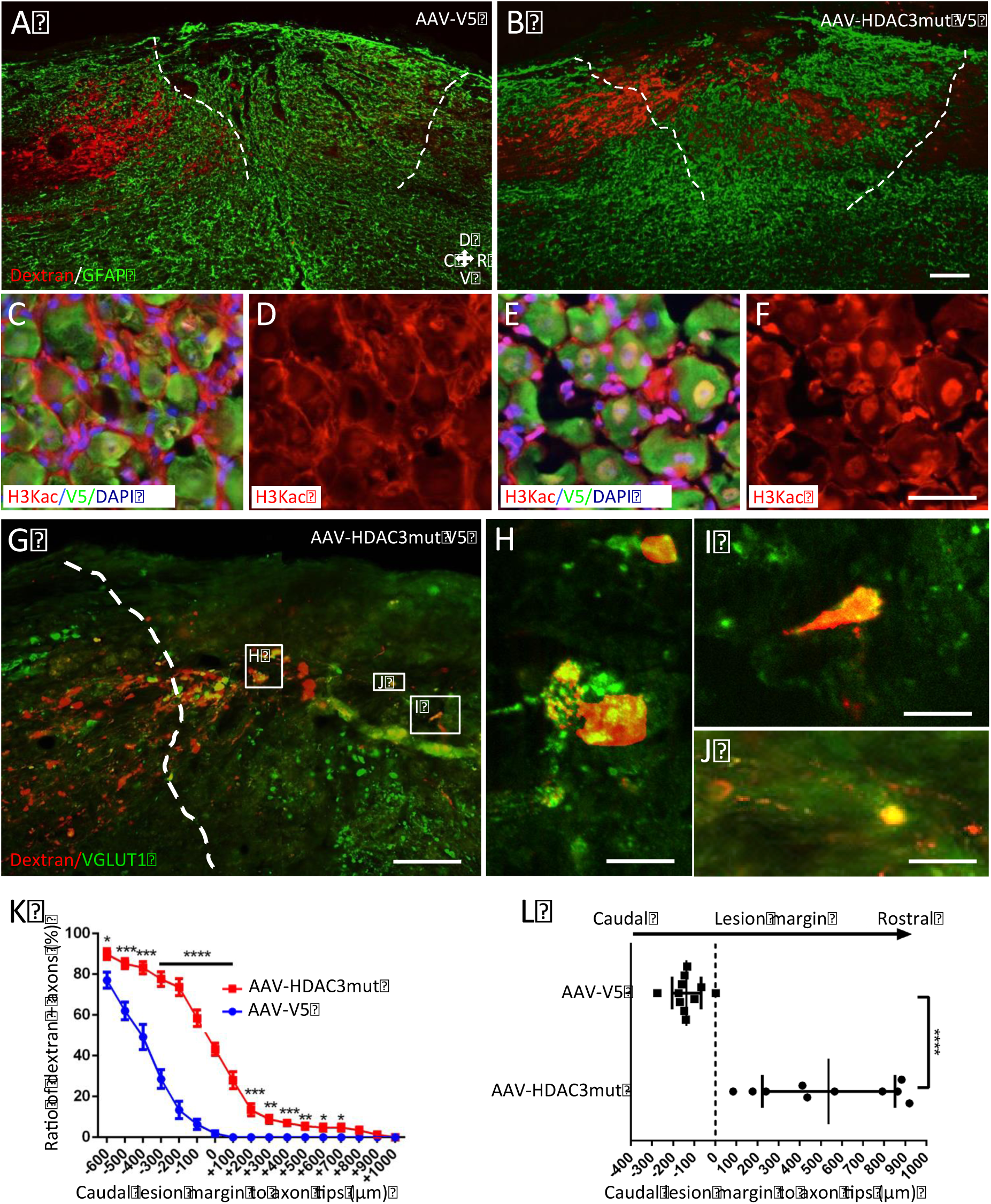
Genetic inactivation of HDAC3 deacetylase activity with HDAC3 dead mutant promotes DRG regenerative growth *in vivo* after SCI. A-F. AAV-HDAC3 mutant (Y298H; dominant negative deacetylase inactive) promoted DRG regenerative growth after spinal cord injury vs AAV-V5 control (injected 5 weeks prior to SCI in the sciatic nerve bilaterally) as shown by dextran-red traced axons across and beyond the lesion site (GFAP, green) (dotted lines denote the rostral and caudal margins of the scar around the lesion). (A-B). AAV-HDAC3 mutant induced upregulation of H3K9ac *in vivo* in DRG neurons, but not in surrounding nuclei of satellite cells (C-F). Scale bar, (A-B) 200 μm, (C-F) 50 μm. G-J. New synaptic formations (VGLUT1+) from regrowing axons are shown beyond the lesion site after injection of AAV-HDAC3mut. Scale bar, (G) 200 μm, (H-J) 50 μm. K-L. Quantification of dextran positive axons shows that AAV-HDAC3mut promotes DRG regenerative growth across and beyond the spinal lesion site. Data is expressed as percentage of dextran+ axons at each distance vs dextran+ axons at −700μm from the lesion margin (K) or distance from the caudal margin of the lesion to the most rostral dextran+ axon tip for each animal (L). N= 10 animals per condition. *p<0.05, **p<0.01, ***p<0.005, ****p<0.001 indicate a significant difference (ANOVA followed by Bonferroni test).

Taken together, these data suggest that inhibition of HDAC3 activity targets multiple signalling networks and pathways as predicted by bioinformatics analysis (Figure 6B, C), which are associated with a regenerative phenotype, likely by overcoming the repressive chromatin environment upon spinal injury.

### Genetic and pharmacological inhibition of HDAC3 promotes DRG outgrowth ex vivo and axonal growth following spinal cord injury

Lastly, we investigated whether repression of HDAC3 activity by *in vivo* genetic or pharmacological inhibition would enhance DRG regenerative growth across the inhibitory spinal cord environment. Therefore, we aimed to inhibit HDAC3 activity in DRG neurons before a spinal cord injury. AAV-HDAC3mut-V5 or an AAV-V5 control virus were injected into the sciatic nerve of adult mice 4 weeks prior to a T9 spinal cord dorsal column crush, which preferentially severs ascending sensory fibers originating from the DRG. Five days before sacrificing the animals, at day 28 post-injury, the fluorescent axonal tracer dextran was injected in the sciatic nerve to examine axonal growth in the spinal cord. An average of 88±1.9% for the control and 85±1.6% for the HDAC3mut of dextran positive cells showed also positive signal for V5 transduction (Supplementary Figure 8A). Data analysis revealed that inhibition of HDAC3 activity with a HDAC3mut promotes significant axonal growth (Figure 7 A, B, K, L, control for spared axons, Supplementary Figure 7B) and it enhances histone acetylation in transduced DRG neurons (Figure 7 C-F, Supplementary Figure 8C). Furthermore, we observed that axons of mice transduced with HDAC3mut express pre-synaptic vGlut1 as shown by dextran/vGlut1 co-labelling (Figure 7 G-J). In support of a specific role for AAV-HDAC3mut in neurite outgrowth of DRG neurons, we found that intrathecal delivery of AAV-HDAC3mut significantly enhanced neurite outgrowth in *ex vivo* cultured DRG neurons to a higher degree than a control virus or a conditioning lesion (Supplementary Figure 9). Taken together, these data show that inhibition of HDAC3 activity can enhance the regenerative potential of DRG neurons. Next, we also tested whether intrathecal administration of *RGFP966* through osmotic minipumps could also enhance growth of DRG sensory axons after spinal cord injury in a model of dorsal hemisection in the adult mouse. *RGFP966* or vehicle was delivered for two weeks post-injury through an intrathecal catheter placed in proximity to the lesion site from the time of the spinal injury. Axonal labelling of dextran traced DRG sensory ascending fibers into and past the lesion was evaluated together with the measurement of the glial scar five weeks after SCI. Importantly, we found that mice treated with RGFP*966* showed increased growth of DRG sensory axons (Figure 8A-D) and as expected, *RGFP966* led to significant enhancements in histone acetylation in DRG (Figure 8E, F). We included in our analysis only spinal cords without sparing of axon fibers as shown by cross sections of cords far rostral to the lesion site (Figure 8G). Importantly, the glial scar and anti-CD11b immunoreactivity around the injury site remained unaffected in RGFP966 versus vehicle treated mice (Supplementary Figure 10A-F). Similarly, intrathecal delivery of the HDAC3 inhibitor *RGFP966* triggered significant neurite outgrowth in ex vivo cultured DRG neurons compared to vehicle on both growth permissive and inhibitory substrates (Supplementary Figure 11).

**Figure 8.**
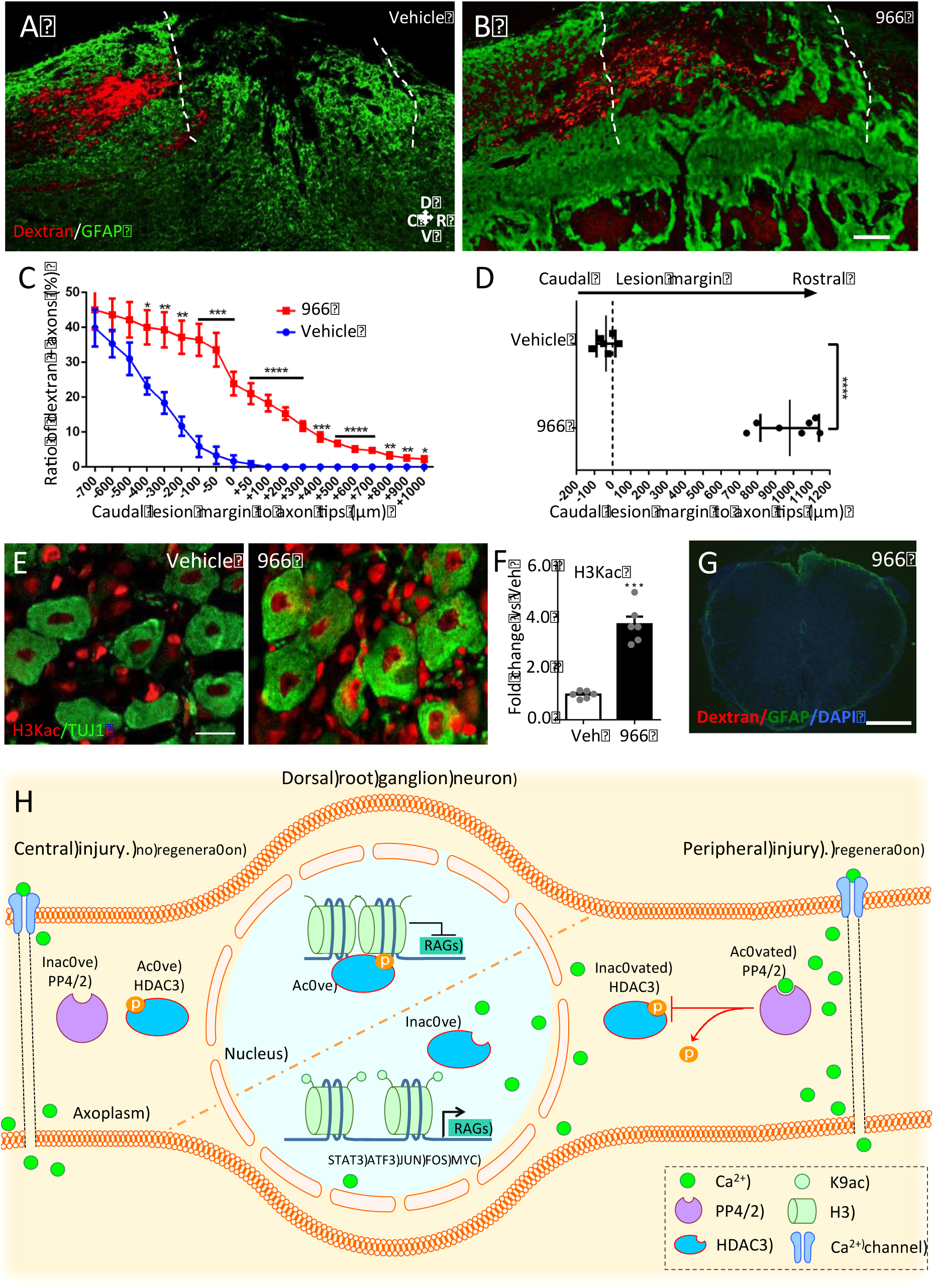
Pharmacological inhibition of HDAC3 promotes DRG regenerative growth. A-B. Intrathecally administered RGFP966 (966) through osmotic minipump for 14 days promoted DRG regenerative axonal growth after spinal cord injury as shown by dextran-red traced axons across and beyond the lesion site (GFAP, green) (dotted lines denote the rostral and caudal margins of the scar around the lesion). Scale bar, 200 μm. C-D. Quantification of axonal regeneration shows dextran axonal labelling across and beyond the spinal lesion site. Data is expressed as ratio of dextran^+^ axons vs dextran^+^ axons at −700 μm from the lesion site ± s.e.m (C) or distance from the caudal margin of the lesion to the last dextran^+^ axon tip (D). N= 10 animals per condition. (*p<0.05, **p<0.01, ***p<0.005, ****p<0.001) indicate a significant difference (ANOVA followed by Bonferroni test). E. Immunofluorescence shows upregulated H3K9ac *in vivo* in DRG neurons after 966. Scale bar, 50 μm. F. Data is expressed as fold change of fluorescence intensity vs veh± s.e.m. N= 6. (***p<0.005) indicate a significant difference (Student’s t-test). G. Spinal cord coronal section of 966 treated mice 10 mm rostral from the lesion core, showing the absence of spared axons. Scale bar, 500 μm. H. Summary diagramme: following central nervous system (CNS) spinal injury, protein phosphatase 4/2 activity is not induced since calcium levels remain unchanged compared to uninjured conditions. HDAC3 remains phosphorylated and occupies deacetylated chromatin contributing to its compaction inhibiting gene expression. Following peripheral nervous system (PNS) sciatic injury, protein phosphatase 4/2 activity is induced by calcium. HDAC3 is dephosphorylated leading to its inhibition and release from chromatin sites contributing to increase in histone acetylation and in the expression of regeneration associated genes (RAGs).

## Discussion

Our data propose calcium-dependent activation of PP4c and to a lesser extent of PP2a leading to dephosphorylation of HDAC3 as a novel molecular mechanism that discriminates between axonal regeneration versus regenerative failure in DRG sensory neurons. Specifically, we found that calcium increases in DRG following a regeneration-competent sciatic but not a regeneration-incompetent spinal injury to activate PP4c that is required for HDAC3 dephosphorylation. While PP4 has been shown to play a role in phagocytic clearance of degenerating axons in Drosophila^21^, its role in axonal regeneration remained unexplored so far. Similarly, while calcium has been linked to the activation of multiple signalling pathways in injured axons, it was previously unknown whether a central spinal lesion associated with regenerative failure would fail to trigger significant increases in calcium in the DRG cell bodies. Here, we found that calcium, which is required for PP4/2 activation is induced in DRG following peripheral regenerative axotomy only.

HDAC3 dephosphorylation inhibits HDAC3 activity following a sciatic nerve injury, while it remains elevated after a non-regenerative central spinal injury. Reduced HDAC3 activity in turn allows for increased histone acetylation and regenerative gene expression in DRG. Indeed, inhibition of PP4/2 activity restricts DRG regenerative growth while inhibition of HDAC3 phosphorylation or activity leads to an enhancement in DRG neurite outgrowth on both growth permissive and inhibitory substrates. Subsequent bioinformatics analysis integrating the HDAC3 protein-protein interaction databases combined with DRG *ex vivo* H3K9ac ChIPseq and RNAseq after a central spinal versus peripheral sciatic lesion further supported a role of HDAC3 in restricting the regeneration programme. Additionally, we found that *in vivo* HDAC3 pharmacological inhibition following spinal cord injury promotes the promoter acetylation and protein expression in DRG neurons of several regeneration-associated genes belonging to signalling pathways that were identified by our combinatorial bioinformatics analysis. They include several transcription factors such as JUN, MYC, STAT3, ATF3 and FOS within key signalling pathways such as MAPK, Insulin and JAKSTAT.

Ultimately, the genetic or pharmacological inhibition of HDAC3 led to significant DRG regenerative growth after spinal cord injury.

While a direct role for HDAC3 in axonal regeneration had not been reported so far, a recent study found that HDAC3 plays also a role in the control of Schwann cell-dependent myelination in the peripheral nerve, where pharmacological inhibition of HDAC3 with RGFP966 promotes remyelination and functional recovery in the injured peripheral nervous system(**21**).

It has also been shown that HDAC3 might control the expression of immune-related gene targets in inflammatory models of SCI such as contusion injuries, where systemic pharmacological HDAC3 inhibition with RGFP966 was followed by improved functional recovery (*22*). However, HDAC3 systemic pharmacological inhibition did not alter the extent of the CD11b positive inflammatory response around the injury site after spinal contusion(*22*), in line with what we found here through intrathecal administration of RGFP966 after spinal dorsal hemisection. In contrast to our findings that intrathecal delivery of RGFP966 promotes DRG regenerative growth in culture, which were supported by enhanced DRG neurite outgrowth following genetic HDAC3 inhibition. The same authors observed that systemic administration of RGFP966 did not affect neurite outgrowth of DRG neurons at 48 hours in culture(*22*). The different route of administration and the timing of when neurite outgrowth has been measured might explain this discrepancy. Importantly, neurite outgrowth of DRG neurons on growth permissive substrates is typically measured within 12-24 hours in culture before they reach an exponential growth state and differences in outgrowth are no longer measurable.

Our data also suggests that axonal regeneration of sensory neurons does not depend upon the generalized inhibition of HDACs or of DNA methyltransferases, as modifying the activity of the various histone-modifying enzymes within our screen had no effect on DRG regenerative growth. This is in line with the poor or absent representation of DNA methylation on gene regulatory regions of DRG after nerve or spinal injury(*15*) and the absent or limited regenerative potential of generalized HDAC inhibition after optic nerve or spinal injury(*12*, *13*). It is however specific signalling pathways that shape the transcriptional environment of DRG neurons that determine whether axonal regeneration succeeds or fails. This will restrict the regrowth programme after spinal cord injury and in contrast, it will work as a central regulatory hub for regenerative signalling following DRG sensory axonal injury. We previously discovered that HATs CBP/p300 and P/CAF form a transcriptional complex with the transcription factor p53 in primary neurons to enhance promoter accessibility of regenerative genes via an active chromatin state on select regenerative gene promoters (*23*-*25*). This was observed both *in vitro* and *in vivo*, where the p53-CBP/p300-P/CAF transcriptional complex was essential for neurite outgrowth and therefore activated during the axonal regenerative programme (*24*, *25*), which is dependent upon an active p53 including after SCI (*26*). In addition, we have recently found that viral overexpression of the histone acetyltransferase p300 can promote axonal regeneration of the optic nerve after optic nerve crush (*12*), while overexpression of P/CAF promotes regeneration after spinal cord injury suggesting that both transcriptional and acetylation-dependent epigenetic mechanisms can directly sustain axonal regeneration in these sensory neurons. Studies by others have recently shown that p300 forms a complex with the transcription factor SMAD1 in DRG after a conditioning lesion (*27*). It is therefore possible that post-injury calcium levels that regulate PP4c activity determine whether active or inactive HDAC3 restricts or allows the accessibility of regenerative transcriptional complexes such as p300 and P/CAF as well as with the associated transcription factors, including SMAD1 and p53 to specific promoters.

Although the axonal injury-dependent regulatory mechanism upon HDAC3 does not involve nuclear-cytoplasmic translocation, but rather PP4 and calcium-dependent HDAC3 dephosphorylation, this is conceptually in line with the reported injury and calcium-depended cytoplasmic export of HDAC5, which enhances axonal regeneration in the PNS(*28*). Interestingly, since HDAC5 does not seem to have catalytic activity(*29*), it is possible that peripheral nerve regeneration may be controlled via interaction between HDAC5 and HDAC3 activity. However, whether HDAC5 plays a role in discriminating between regenerative success or failure such as following spinal cord injury was not investigated.

Since whole DRGs contain both neurons and satellite cells, the development of combinatorial single cell RNAseq and ChIPseq will hopefully soon allow studying genetic and epigenetic changes specifically in neuronal populations, adding further specificity to unbiased molecular screenings. However, immune cell infiltration into the ganglion induced by a peripheral nerve typically occurs at a much later time points (7 to 14 days after lesion)(*30*, *31*) than the one used in this study (24h), where we did not observe any accumulation of inflammatory cells in sciatic ganglia.

Taken together our findings show a novel mechanism for the regulation of the differential regenerative ability between regeneration-competent peripheral versus regeneration-incompetent central axonal injury. We found that (i) calcium-dependent activation of PP4 leads to HDAC3 dephosphorylation causing in turn HDAC3 enzymatic inhibition; (ii) HDAC3 restricts the axonal regeneration programme; (iii) pharmacological or genetic inhibition of HDAC3 activity promotes regenerative signalling and (iv) enhances the axonal growth state of sensory neurons partially overcoming regenerative failure after SCI.

## Acknowledgments

We would like to thank start up funds-Division of Brain Sciences, Imperial College London (SDG), and the Hertie Foundation for financial support (SDG); Wings for Life (SDG); the DFG (SDG), the Henry Smith Charity and Rosetrees Trust (SDG). The research was supported by the National Institute for Health Research (NIHR) Imperial Biomedical Research Centre (SDG). The views expressed are those of the author(s) and not necessarily those of the NHS, the NIHR or the Department of Health.

## Materials and Methods

### Mice

Animal work was carried out in accordance to regulations of the UK Home Office. Wild-type C57Bl6/J (Harlan) mice ranging from 6 to 8 weeks of age were used for all experiments. Male and female were used in a 50-50% basis for each study. Each animal was randomly assigned to an experimental group. For all surgeries mice were anesthetized with isofluorane (5% induction 2% maintenance) and a mixture of buprenorphine (0.1mg/kg) and carprophen (5mg/kg) was administered peri-operatively as analgesic. Animals outside the 6-8 week or 15-30gr range were excluded for the use in any study. Animals wounded or with clear signs of distress were also excluded. All animal procedures were approved by Imperial College London ethic committee, and were performed in accordance with the UK Animals Scientific Procedures Act (1986).

### Compounds

RGFP233, 933 and 966 were received from Repligen. GW376713X (13X), GSK1786269A (69A), GSK1210151A (51A), GSK343A (43A), GSK2961917A (17A), GSK2668977A (J4), GSK2801 (801), GSK484 (484), GSK2924467A (67A), GSK2980071B (71B) were obtained from a collaboration through an MTA with GSK. EGTA and KCl were purchased from Sigma and Fostriecin from Cayman. For each group treated with a drug, the respective control group received the same volume of vehicle.

### Dorsal column axotomy (DCA)

Surgeries were performed as previously reported (*26*). Briefly, mice were anesthetized and a T9 laminectomy was performed (∽20mm from the sciatic DRGs), the dura mater was removed, taking care of not damaging the spinal cord. A dorsal hemisection until the central canal was performed with fine forceps (FST). For the control laminectomy surgery, the dura mater was removed but the dorsal hemisection was not performed.

### Sciatic nerve axotomy (SNA)

Briefly, sciatic nerve lesion experiments were performed under isoflurane anesthesia. The biceps femoris and the gluteus superficialis were separated by blunt dissection, and sciatic nerve was exposed. Sciatic injury was performed by a sharp axotomy with iridectomy scissors (FST). Sham-operated mice that underwent exposure of the sciatic nerve without axotomy were used as surgery controls.

### Dorsal Root Ganglia (DRG) cell culture

Adult DRGs were dissected and collected in Hank’s balanced salt solution (HBSS) on ice. DRGs were transferred to a digestion solution (5mg/ml Dispase II (Sigma), 2.5mg/ml Collagenase Type II (Worthington) in DMEM (Invitrogen) and incubated at 37°C for 45 mins with occasional mixing. Thereafter DRGs were transferred to media containing 10% heat inactivated FBS (Invitrogen), 1X B27 (Invitrogen) in DMEM:F12 (Invitrogen) mix and were manually dissociated by pipetting until no remaining clumps of DRGs were observed. Next, single cells were spun down, resuspended in media containing 1X B27 and Penicillin/Streptomycin in DMEM:F12 mix and plated at 3500/coverslip. The culture was maintained in a humidified atmosphere of 5% CO_2_ in air at 37°C. Cells were allowed to grow for 12h in culture before fixation.

### Ex-vivo DRG cell culture

Animals underwent intrathecal injections of 5μL RGFP966 (10 or 100μM), Fostriecin (240μM) or vehicle (5% DMSO in 0.9% saline) or 2μL of AAV1 (control or HDAC3mut) into Cerebrospinal fluid (CSF) between L5 and L6 laminae, or alternatively 100 μL of EGTA 10mM or vehicle (0.9% saline) were applied to sciatic nerve before performing sham or SNA surgeries as above mentioned. 24hr later sciatic bilateral DRGs were dissociated and plated following the above protocol.

### Calcium imaging

DRG primary cells were cultured for 12h following standard protocol. Cells were washed with Ringer solution and loaded with Fluo4AM (2 μM, Thermo Scientific) diluted in Ringer solution, after 30 min incubation at RT protected from light. Cells were washed, and KCl-Ringer solution was applied (40 mM KCl). For Fostriecin treated cultures, cells were pretreated for 30 min with vehicle (1%DMSO) or Fostriecin (200nM) before KCl administration. Cells were monitored for KCl-induced Fluo4-AM fluorescence increases. Cells were then fixed, such that the KCl-induced increase in fluorescence was frozen, and subsequently stained for H3K9ac, pHDAC3 or HDAC3. Correlation between Fluo4AM and immunostainings levels were quantified and plotted.

### PP4/PP2A activity

Protein phosphatase 4c and 2A activities were measured by measuring phosphatase activity (PP2A phosphatase activity kit, Millipore, CA, USA) on DRGs after central and peripheral injury prior PP4 or PP2A immunoprecipitation. Briefly, sciatic DRGs were extracted 24h after surgery, protein lysates were obtained with 1 liter of imidazole buffer (20 mM imidazole-HCl, 2 mM EDTA, 2 mM EGTA, pH 7.0 with 10 μg/mL each of aprotinin leupeptin, pepstatin, 1 mM benzamidine, and 1 mM PMSF) per 25 g of tissue, homogenized with micropestles on ice and centrifuged at 2000 x g for 5 minutes at 4°C. Protein lysate concentration was measured by BCA, and 100μg of protein were used for each reaction, together with 4μg of PP2A (Millipore, CA, USA), PP4 (PPX, Santa Cruz, CA, USA) antibodies or Mouse IgG (Millipore, CA, USA). Samples were processed following the kit instructions and activity of samples was then measured in an Infinite M200Pro microplate reader (Tecan) at 650nm according to manufacturers guidelines.

### *Ex vivo* calcium assay

DRG were dissected 24h following sham, SNA, laminectomy or DCA. Samples homogenates were employed for calcium measurement by using the Calcium Detection Assay Kit (Abcam). In particular, sciatic DRG from 3 mice per replicate were homogenized with a sonicator in 120 μL Calcium Assay Buffer on ice and the collected supernatant was used for calcium detection on a microplate reader (OD575 nm) following the manufacturer’s protocol. Calcium concentration was calculated in 4 biological replicates for each condition.

### PP4/PP2A RNA silencing

Following DRG dissection and dissociation, cells were washed twice with HBSS and transfected with a mixture of a GFP and siRNA plasmids mixture using Amaxa 4D-Nucleofector kit (Lonza). Briefly, 50,000 DRG cells per reaction were resuspended with 20 μL Nucleocuvette Strip (16.4 μL Nucleofector solution, 3.6 μL Supplement) and 0.4 μg pmaxGFP mixed with 6 pmol of control, PP2a or PP4c siRNAs (Santa Cruz) were applied per reaction. After nucleofection, cuvettes were incubated 10 min at RT and cells were resuspended with pre-warmed DRG medium. 25,000 cells were plated per coverslip for 72h. Due to the lack of efficient antibodies against PP4, verification of silencing was performed via immunocytochemistry as decribed below and PCR using the primers provided in the kit (Santa Cruz, sc-39203-PR). For PP2a silencing, verification of silencing was performed via immunoblotting as described below.

### Mutant HDAC3 variants transfection in DRG cultures

HDAC3 constructs were cloned on a pAAVIREShrGFP backbone (Agilent). FLAG tag, the IRES element and the GFP were removed from the original backbone via restriction on the BamHI site in the MCS and the second Kpn1 site after GFP, and was replaced with either the V5 tag MW87 (empty vector), HDAC3 (seq. accession number NM_01411)+V5 tag (wildtype HDAC3), HDAC3 S424A+V5 tag, HDAC3 S424D+V5 tag, HDAC3 Y298H+V5 tag (HDAC3mut). Briefly, DRGs were dissected and dissociated as mentioned previously, cells pellets were resuspended after digestion with an OptiMEM (Invitrogen) solution containing 10μl/ml of Lipofectamine2000 (Invitrogen) and 6.25μg/ml of DNA. Cells were allowed 3h for the transfection, and then transfection solution was replaced with normal DRG medium. Cells were fixed after 12h.

### RNA sequencing

Sciatic DRGs (3 biological replicates, pool of 2 mice/replicate) were extracted 24h after sham, sciatic nerve axotomy, laminectomy or dorsal column axotomy surgeries (surgeries were performed as described above), and RNAseq was performed and analysed as we previously described^20^. Sequence reads were aligned to the mm10 mouse reference genome sequence using tophat version 2.0.12 running Bowtie2-2.2.3. Gene structure annotations corresponding to the Ensembl annotation of the mm10 genome sequence were used to build a transcriptome index and provided to tophat during the alignment step. The aligned reads were sorted using samtools-0.1.19, and then read counts per gene were obtained from mapped reads using HTSeq-0.6.1. EdgeR version 3.8.6 (using limma-3.22.7) in R-3.1.1 was used to identify differentially expressed genes. Following the procedures described in the edgeR documentation, read count tables were loaded into R. GC-content bias was controlled for using full quantile normalization in EDASeq-2.0.0 running in R-3.1.1. EDASeq was also used to generate quality-checking figures before and after this normalization. Differential expression testing was performed on the normalized output from EDASeq using edgeR. Read-level quality checking was performed using fastqc-0.10.1 and the fastqc-aggregator (https://github.com/staciaw/fastqc_aggregator) and gene-level quality checking was performed using RSeQC-2.6.1. The cut-off criteria for the differentially expressed (DE) genes was set at FDR< 0.05.

### H3K9ac ChIP sequencing

Sciatic DRGs (2 biological replicates, pool of 20 mice/replicate) were extracted as above, 24 hours after sham, sciatic nerve axotomy, laminectomy or dorsal column axotomy surgeries. Chromatin IP was performed according to our previously published protocol^14^ with a few adjustments. Briefly, DRG pellets were crushed using an automatic pestle and chemically cross-linked with a 1% formaldehyde solution containing 50mM Hepes-KOH pH7.5, 100mM NaCl, 1mM EDTA and 0.5mM EDTA, for 15 minutes at room temperature. Nuclear extracts were then sonicated for 30 min using a Bioruptor (Diagenode) and successful chromatin shearing (200-800 bp) was confirmed by agarose gel analysis. Immunoprecipitation was performed overnight with Protein G Dynabeads (Invitrogen) bound to 10 μg of H3K9ac antibody (Ab10812, Abcam). After washing, elution and reverse crosslinking, DNA was treated with RNAse A and Proteinase K and purified using Qiagen PCR purification columns. The concentration of the recovered DNA was quantified with Picogreen assay and the enrichment of the immunoprecipitate calculated with respect to the input and IgG. Library preparation was performed using the NEBNext Ultra DNA Library Prep Kit from Illumina (NEW ENGLAND BioLabs) following the manufacturer’s protocol. Briefly, 30 ng of Input and IP DNA were end-repaired and adaptor ligated, using a dilution of 1:10 of adaptors. Samples were cleaned to remove unligated adaptors and size selection was performed to select and enrich fragments of 200bp in size. Libraries were amplified by PCR using multiplex oligo following manufacturer’s protocol. Prior to sequencing, libraries were run on an Agilent Bioanalyser for size and quality control checking. Sequencing was performed in an Illumina HiSeq2000, generating 50bp single ended reads. Input libraries were used as background references for peak identification, sequence alignment, differential expression analysis and identification of H3K9ac-associated peaks.

For ChIPseq analysis, transcripts were aligned to the mm10 reference genome using Bowtie2-2.2.3, sorting reads in Samtools-0.1.19. Genomic bins of 1000bp upstream and downstream of each transcription start site for each gene were created using the same gene annotation as used for the RNAseq data. Read counts per genomic bin were obtained from the mapped reads using HTSeq-0.6.1 and subsequently, differential binding testing was conducted in EdgeR-3.8.6. Quality examination on aligned reads was performed using ChIPQC-1.2.2. The cut-off criteria for the differentially occupied genes by H3K9ac was set at *p* value <0.05. ChIP signal distribution plots were generated using NGSplot version 2.47.1 Briefly, the ChIP signal for each condition is plotted along a gene body region either for all genes in the annotation or a select subset of genes. The expression and differential expression data from the RNAseq experiment were used to select the groups of genes to sample the ChIP signal in order to correlate the ChIP signal with changes in the gene’s expression between biological conditions. The ChIPseq signal tracks were generated using the macs2 bdgcmp command. The signal pileups of the ChIP and input conditions produced by macs2 callpeaks command were used as the treatment and control inputs, respectively, for the bdgcmp command. The output bedgraph reports the –log10(pvalue) of treatment relative to control for each bin using the ‘–m ppois’ option. The bedgraph files were then converted to bigwig files using the bedGraphToBigWig utility from KentUtils. The RNAseq signal tracks were created using the deeptools utility, bamCoverage. The bam files of each replicate per condition were pooled together for use as input to bamCoverage (*32*).

## Data availability

RNAseq and H3K9ac ChIPseq data have been deposited at the Gene Expression Omnibus (GEO) with accession codes: GSE97090 for RNAseq, https://www.ncbi.nlm.nih.gov/geo/query/acc.cgi?token=anmdoqiodzgfvat&acc=GSE97090) and GSE108806 (exwheyyitzqlpoz) for ChIPseq.

### HDAC3 Network

The signalling network was generated using the protein interactome (calculated with Fpclass, score ≥ 0.8) of HDAC3 and H3K9ac dependent genes (genes upregulated (*p* value <0.05) following SNA with an increased H3K9ac occupancy (*p* value <0.05). Only upregulated proteins after SNA were used to build the network. Any nodes in which no connections were found were removed from the output. The network was visualized using Cytoscape software (version 3.3.0), circularizing the terms that have been found enriched for signalling pathways in a KEGG enrichment analysis (*p* value <0.05, see Supplementary File 5).

### Immunoblotting

Proteins from either sciatic DRGs or Dissociated DRG neurons were extracted using RIPA buffer with protease and phosphatase inhibitor cocktails (Roche). Total lysates were obtained by 30’ centrifugation at 4°C. Nuclear-/cytoplasmic-protein lysates from DRGs were extracted using the NE-PER Nuclear and Cytoplasmic Extraction Reagents (Thermo Scientific) following manufacturer’s instruction. Protein concentration of lysate was quantified using Pierce BCA Protein Assay Kit (Thermo Scientific). 10-50μg of proteins were loaded to SDS-PAGE gels and transferred with iBlot Dry Blotting System (Thermo Scientific). Membranes were blocked with 5% BSA or milk for 1h RT and incubated with HDAC3 (1:1000, Abcam), pHDAC3 (1:1000, Cell Signaling Technology (CST)), H3K9ac (1:1000, CST), H3 (1:1000, CST), H3 (1:1000, CST), GAPDH (1:1000, CST), ERK (1:1000, CST), pERK (1:1000, CST) or PP2A (1:500, Santa Cruz) at 4°C O/N. Following HRP-linked secondary antibody (GE Healthcare) incubation for 1h RT, membranes were developed with ECL substrate (Thermo Scientific).

### Immunocytochemistry (ICC)

Glass coverslips were coated with 0.1mg/ml PDL, washed and coated with mouse Laminin 2ug/ml (Millipore). For myelin experiments, they were additionally coated with 4μg/cm^2^ rat myelin. Cells were plated on coated coverslips for 12h, at which time they were fixed with 4% paraformaldehyde/4% sucrose. Immunocytochemistry was performed by incubating fixed cells with anti-αlll Tubulin (Tuj-1, 1:1000, Promega), GFP: (1:1000, Abcam), HDAC3 (1:1000, Abcam), pHDAC3 (1:200, CST), H3K9ac (1:1000, CST), PP4c (1:200, Santa Cruz) or anti-V5 (1:200, Millipore) antibodies at 4°C O/N. This was followed by incubation with Alexa Fluor 564 or 488 conjugated goat secondary antibodies according to standard protocol (Invitrogen). All cells were counterstained with Hoechst (Molecular Probes).

### Immunohistochemistry (IHC)

Dissected DRG were fixed in 4% PFA and transferred to 30% sucrose at 4°C overnight. Tissues were embedded in OCT compound (Tissue-Tek), frozen and cryosectioned. 10 μm tissue sections were permeabilized and blocked in 10% NGS, 0.3% Triton X-100, PBST for 1h at RT followed by primary antibodies incubation with βIII tubulin (Tuj-1, 1:1,000, Promega), GFP: (1:1000, Abcam), HDAC3 (1:1000, Abcam), pHDAC3 (1:200, CST), H3K9ac (1:1000, CST), anti-V5 (1:200, Millipore), p-Stat3 (1:100, CST, #9145), c-Jun (1:50, CST,#9165), ATF3 (1:100, Santa Cruz, sc-188), p-Erk (1:250, CST, #9101), IGF1R (1:200, CST, #3027), Myc (1:100, Sigma, M4439), NF200 (Rabbit, 1:80, Sigma, N4142; mouse, 1:400, Sigma, N0142) at 4°C O/N. This was followed by incubation with Alexa Fluor 564 or 488 conjugated goat secondary antibodies according to standard protocol (Invitrogen). All sections were counterstained with Hoechst (Molecular Probes).

### Image Analysis for IHC and ICC

DRG or spinal cord were taken at 20X magnification with an Axioplan 2 (Zeiss) microscope and processed with the software AxioVision (Zeiss). Exposure time and gain were maintained constant between conditions for each fluorescence channel. A modified protocol based on the guidelines described on (Protocol Exchange (2012) doi:10.1038/protex.2012.008) was used to quantify fluorescence intensity. Briefly, using Image J, a constant fluorescence intensity threshold was set across samples. Based on the threshold, for each picture the intensity density (ID) of pixels were calculated in each channel and then divided by its respective number of cells (about 225 cells/picture). This was done in triplicate and blind to the experimental group.

### Neurite Length Analysis

Immunofluorescence was detected using an Axiovert 200 microscope (Zeiss) and pictures were taken as a mosaic at 10X magnification using a CDD camera (Axiocam MRm, Zeiss). 5 fields per coverslip were included in the analysis. All analyses were performed in blind. Neurite analysis and measurements were performed using Neurite J plugin for Image J software (Image J). Approximately between 30 and 50 cells were analyzed per each condition. Total neurite length per neuron was quantified and averaged (12 hours in culture).

### HDAC3 activity

HDAC3 activity was measured by measuring HDAC activity on DRGs after central and peripheral injury prior HDAC3 immunoprecipitation. Briefly, sciatic DRGs from 3 animals were extracted 24h after surgery and pooled per sample, protein lysates were obtained using non denaturizing lysis buffer (NDLB, 20 mM Tris-HCl pH 8, 137 mM NaCl, 1% IGEPAL, 2 mM EDTA), for the HDAC3 immunoprecipitation total lysates were bound to HDAC3 antibody (rabbit, Abcam), pulled down with Protein G magnetic beads, washed with NDLB, and eluted with 0,1M glycine pH 2, and neutralized with Tris pH 8. Activity of samples of 20μg of immunoprecipitated HDAC3 were measured with fluorometric histone deacetylase activity kit (Active motif, Rixensart, Belgium) according to manufacturers guidelines.

### Intrathecal catheter and osmotic minipump implantation

For RGFP966 delivery, osmotic minipumps (Alzet 2002) were used. A customized 32G i.t. catheter (ReCathCo) was placed subdurally from T11 to T9 caudally connecting osmotic minipump with vehicle or RGFP966 placed in subcutaneous space on the back of mouse. The whole instrument was fixed with cyanoacrylate (Cyano Veneer, Hager&Werken) on muscle-cleaned T12 bone. All pumps were removed at 14 days after injury. Immediately after pump implantation, a T9-T10 laminectomy was performed and a dorsal overhemisection was performed with a microblade (Fine Science Tools).

### In vivo AAV-V5/mutant HDAC3 injection

Briefly, mice were anesthetized with isoflurane inhalation and sciatic nerves were exposed bilaterally. 1.5-2 μL of AAV1-virus were injected with glass-pulled capillary. A dorsal column crush (30 sec) injuring mainly ascending sensory tracts from dorsal columns was performed with a fine #5 microsurgery forceps (Fine Science Tools) two weeks after viral injection.

### Tracing and tissue processing of injured spinal cords

Five weeks after SCI, 2 μL of tracer (15 % Dextran Alexa Fluor 555, 10000MW, Thermo Scientific) were injected into sciatic nerve bilaterally. Spinal cords were dissected 5 days after tracing following transcardial perfusion with PBS and 4% PFA, and subsequently cryopreserved with 30% sucrose overnight at 4°C. Spinal cords and brain stems (Raphe nucleus) were mounted and frozen in OCT mounting media and sectioned at 18 μm on a cryostat. Subsequent GFAP (1:500, Millipore), CD11b (1:1000, Millipore), or VGLUT1 (1:500, Synaptic Systems) stainings were performed as previously described. The GFAP^+^ scar and the CD11b^+^ immunoreactivity within the GFAP positive area were quantified by calculating pixel intensity. A high-resolution image was obtained at 40X magnification using the Zeiss Axioplan Microscope (Axiovert 200, Zeiss Inc.). Images for the same antigen groups were processed with the same exposure time. Assessment of fluorescence intensity was performed using AlphaEaseFC 4.0.1 software by measuring the intensities specifically within the borders of GFAP + scar. At least 3 sections per sample were quantified. The intensity values of each cell were normalized to the background fluorescent signal and mean values of intensities were calculated for each animal.

### Quantification of axonal regeneration

For each spinal cord after dorsal column injury, the number of fibers rostral to the lesion and their distance from the lesion epicenter was analyzed in 4-6 sections per animal with a fluorescence Axioplan 2 (Zeiss) microscope and with the software Stereo-Investigator 7 (MBF bioscience). The lesion epicenter (GFAP) was identified in each section at 40X magnification. The total number of labeled axons rostral to the lesion site was normalized to the total number of labeled axons caudal to the lesion site counted in all the analyzed sections for each animal, obtaining an inter-animal comparable ratio. Sprouts and regrowing fibers were defined following the anatomical criteria reported by Steward and colleagues (*33*).

### HDAC3 RNA silencing

For HDAC3 RNA silencing, shRNA against mouse HDAC3 and scramble shRNA were designed using BLOCK-IT RNAi Designer tool (Life Technologies) and cloned into pRNAT-U6.1/Neo (Gene Script) that carries GFP marker, allowing tracking of the transfected cells. The sequences of the shRNAs were as follows: sense:GGATCCC(GGGTGCTCTACATTGATATCG)TTGATATCCG(CGATATCAATGTAGAGCACCC)TTTTTTCCAAAAGCTT; antisense:GGATCCC(GCCGCTACTATTGTCTCAATG)TTGATATCCG(CATTGAGACAATAGTAGCGGC)TTTTTTCCAAAAGCTT; sense:GGATCCC(GACAGGTCTCTGGAACGTTAT)TTGATATCCG(ATAACGTTCCAGAGACCTGTC)TTTTTTCCAAAAGCTT; antisense:GGATCCC(ATAGTTGTACTCGCGTACGGT)TTGATATCCG(ACCGT ACGCGAGTACAACTAT)TTTTTTCCAAAAGCTT, where the sense and antisense strand, separated by a Loop, indicated between brackets. Briefly, 25000-30000 DRG/transfection were washed 3 times after dissociation with PBS w/o Ca^2+^/Mg^2+^ and transfected using the Neon Transfection System (Life Technologies). Cells were resuspended in 11 microliters of buffer R and transfected with 1 microgram of DNA at 1200 mV, 2 pulses, 20 msec. After recovery, cells were plated in medium w/o antibiotics and after 48h in culture were fixed and analysed.

### H3K9ac ChIP-PCR

Lumbar L1-L6 DRG’s were extracted as described above, 24 hours after dorsal column axotomy surgeries plus intrathecal injection of 5μl of RGFP966 or vehicle (5% DMSO in 0.9% saline). Tissues from 5 animals were pooled per sample. Briefly, DRG pellets were crushed using a pestle and chemically cross-linked with a 1% formaldehyde solution containing 50mM Hepes-KOH pH7.5, 100mM NaCl, 1mM EDTA and 0.5mM EDTA, for 15 minutes at room temperature. Nuclear extracts were then sonicated for 10 min by using a tip sonicator, and successful chromatin shearing (200-800 bp) was confirmed by agarose gel analysis. Immunoprecipitation was performed overnight with Protein G Dynabeads (Invitrogen) bound to 5 micrograms of H3K9ac (Abcam 10812) or Rabbit IgG. After washing, elution and reverse crosslinking, DNA was treated with RNAse A and Proteinase K and purified using Qiagen PCR purification columns. Real-time qPCR was run LightCycler 480 SYBR Green I Master (Roche) in a StepOnePlus cycler (Applied Biosciences). Cts were calculated following the manufacturer’s instructions. Expression values are expressed as 2^−^△△Ct. First ChIP C_t_s were normalized versus IgG to exclude non-specificity (△Ct), then the amount of chromatin was normalized using △Ct from the input. Primers were designed based on H3K9ac enrichment after H3K9ac ChIPseq. Primer sequences used were as follows:

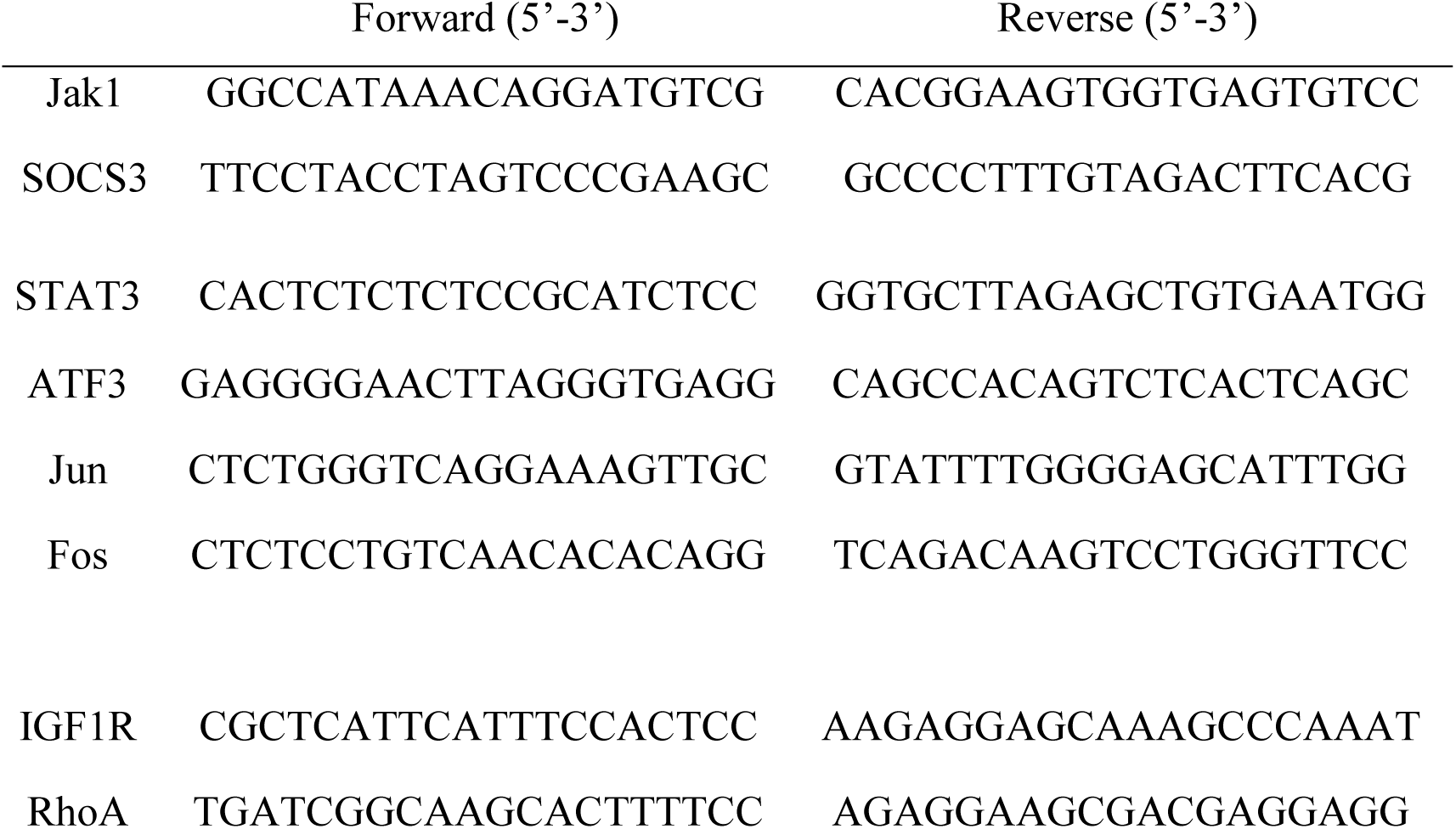

### Analysis of the HDAC3 interactome

To identify how HDAC3 may interact and function at the protein level, an *in silico* analysis was performed using FPClass prediction software to predict high-confidence protein-protein interactions. Interactions were considered significant when the total score was ≥0.4.

### Transcription factor binding site analysis

In order to identify common regulators of the genes that were upregulated in association with increased H3K9ac occupancy, transcription factor binding site enrichment analysis was performed using Pscan software(*34*). Predicted transcription factor binding motifs were screened within the JASPAR 2016 database and enrichment of overrepresented consensus sequences in the promoter region of input genes (−450bp to +50bp) was computed against the *mus musculus* background dataset. Motifs were considered significant when Bonferroni *p* value<0.05. Predicted TFs that could bind to HDAC3 were identified through integration with FpClass HDAC3 predicted PPI (total score ≥0.4).

### Gene Ontology and Pathway analysis

Gene Onytology (GO) and KEGG pathway analysis was performed in DAVID (https://david.ncifcrf.gov/). Enrichment terms were considered significant when *p* value<0.05.

### Quantitative Real-time PCR

Total RNA from bilateral sciatic DRG was extracted 24h after peripheral or central injuries as described above by using RNeasy kit (Qiagen) according to manufacturer’s guidelines. cDNA was then synthesized with SuperScriptTM II reverse transcriptase (Invitrogen). Real-time qPCR was run with KAPA SYBR FAST qPCR kit (Kapa Biosystems) in a 7500 light cycler (Applied Biosciences). Cts were calculated following the manufacturer’s instructions. Expression values are expressed as 2^−△△Ct^. First C_t_s were normalized versus GAPDH as a housekeeping gene, and t the relative amount was normalized against corresponding sham controls. Primer sequences used were as follows:

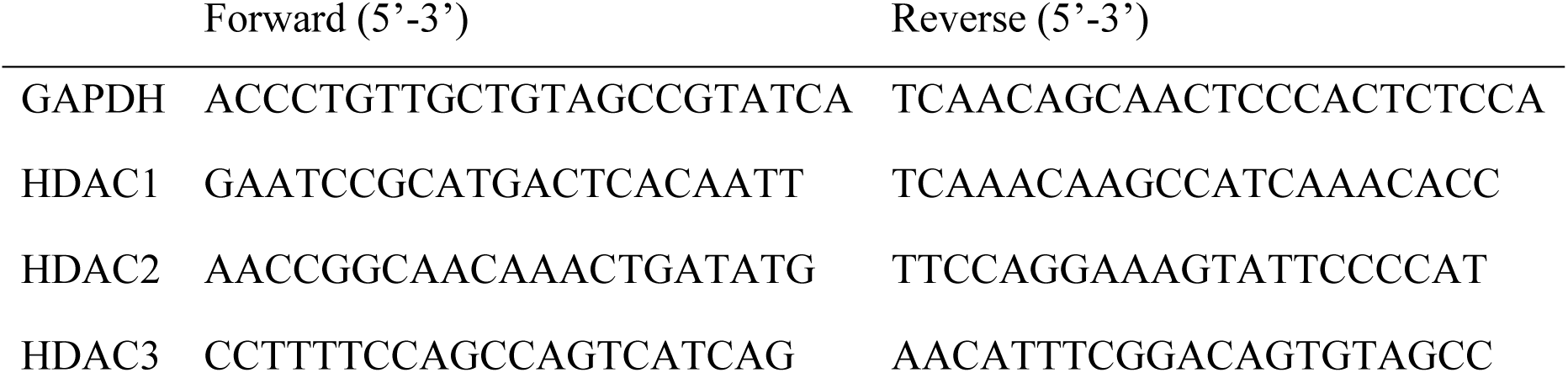

### Statistical analysis

Unless otherwise stated, data is plotted as the mean ± SEM. All experiments were performed three times unless specified. Normality of the distributions was checked via Shapiro-Wilk test, asterisks indicate a significant difference analyzed by ANOVA with Bonferroni post-hoc test or Student’s i-test as indicated (* *p*<0.05; ** *p*<0.01; *** *p*<0.005; **** p<0.001). All tests performed were two-sided, and adjustments for multiple comparisons and/or significantly different variances (Fisher’s F) were applied were indicated. All data analysis was performed blind to the experimental group.

Unless otherwise stated, sample size was chosen in order to ensure a power of at least 0.8, with a type I error threshold of 0.05, in view of the minimum effect size that was expected.

---

**Supplementary Figure 1.**
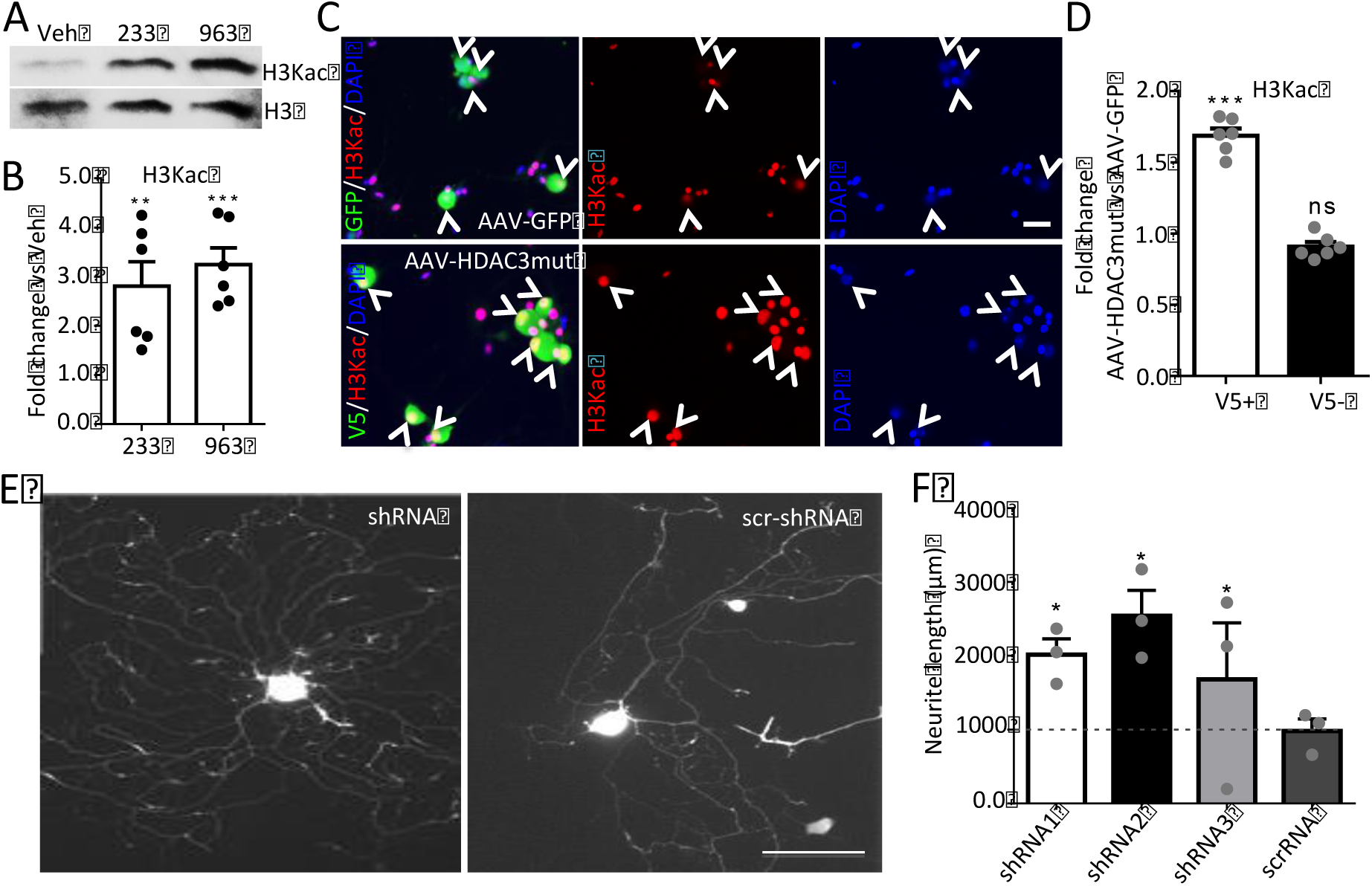
Genetic silencing or inhibition of HDAC3 promotes neurite outgrowth of DRG neurons *in vitro*. A-B. Immunoblot showing the specific increase of H3K9ac after HDAC1 (233, 100nM) or HDAC1/2 (963, 100nM) inhibition on DRGs. B. Data is expressed as mean fold change of immunoblot band intensity ± s.e.m. N= 3. C-D. Immunofluorescence anti-V5 or GFP and H3K9ac after DRG infection with AAV-HDAC3mut-V5 (Y298H) shows increase in H3K9ac as compared to control AAV-GFP (arrowheads). D. Data is expressed as mean fold change vs control AAV-GFP of fluorescent intensity of H3K9ac in V5/GFP positive or negative cells ± s.e.m. N= 3. E. Immunofluorescence anti-TUJ1 following shRNA-mediated HDAC3 silencing vs scrambled (scr-shRNA) in cultured DRG cells shows an increase in neurite outgrowth. Scale bar, 100 μm. F. Data is expressed as mean ± s.e.m. N= 6 (3 independent experiments from 2 biological replicates). (*p<0.05) indicate significant difference versus scrambled shRNA (ANOVA followed by Bonferroni test).

**Supplementary Figure 2.**
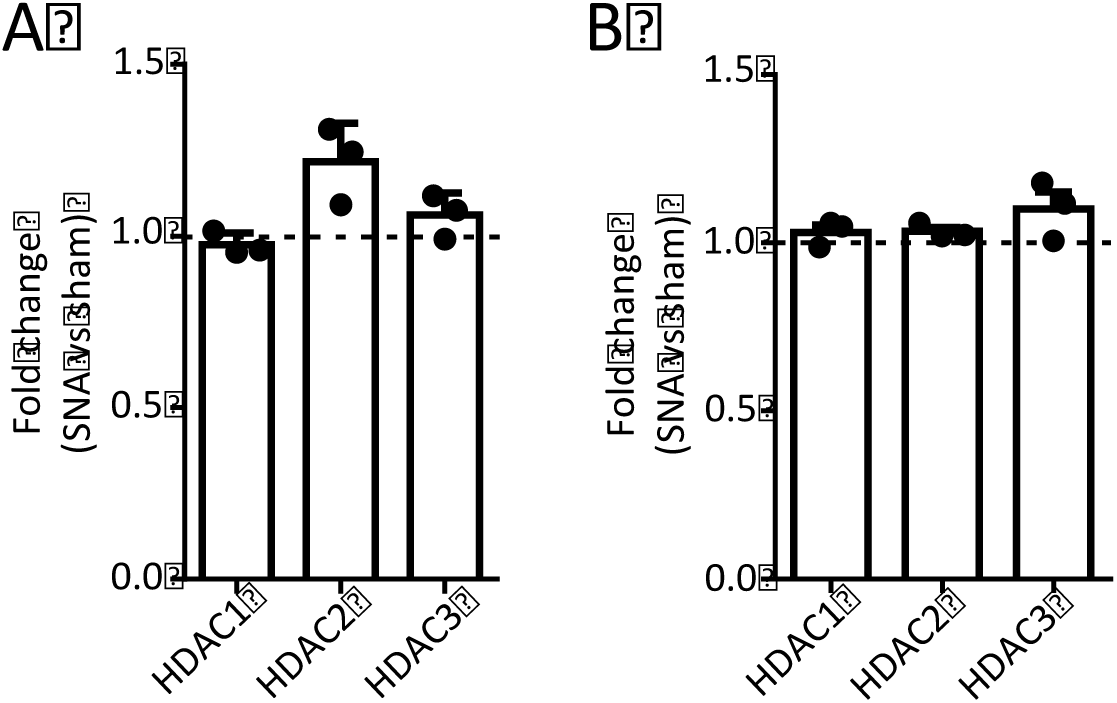
Gene expression of class I HDACs in DRGs after peripheral and central axonal injuries. A-B. RNA levels of HDAC1, 2 and 3 after SNA (A) or DCA (B) vs respective sham or laminectomy. Data is expressed as mean relative expression vs sham or lam ± s.e.m. N= 3 (ANOVA followed by Bonferroni test).

**Supplementary Figure 3.**
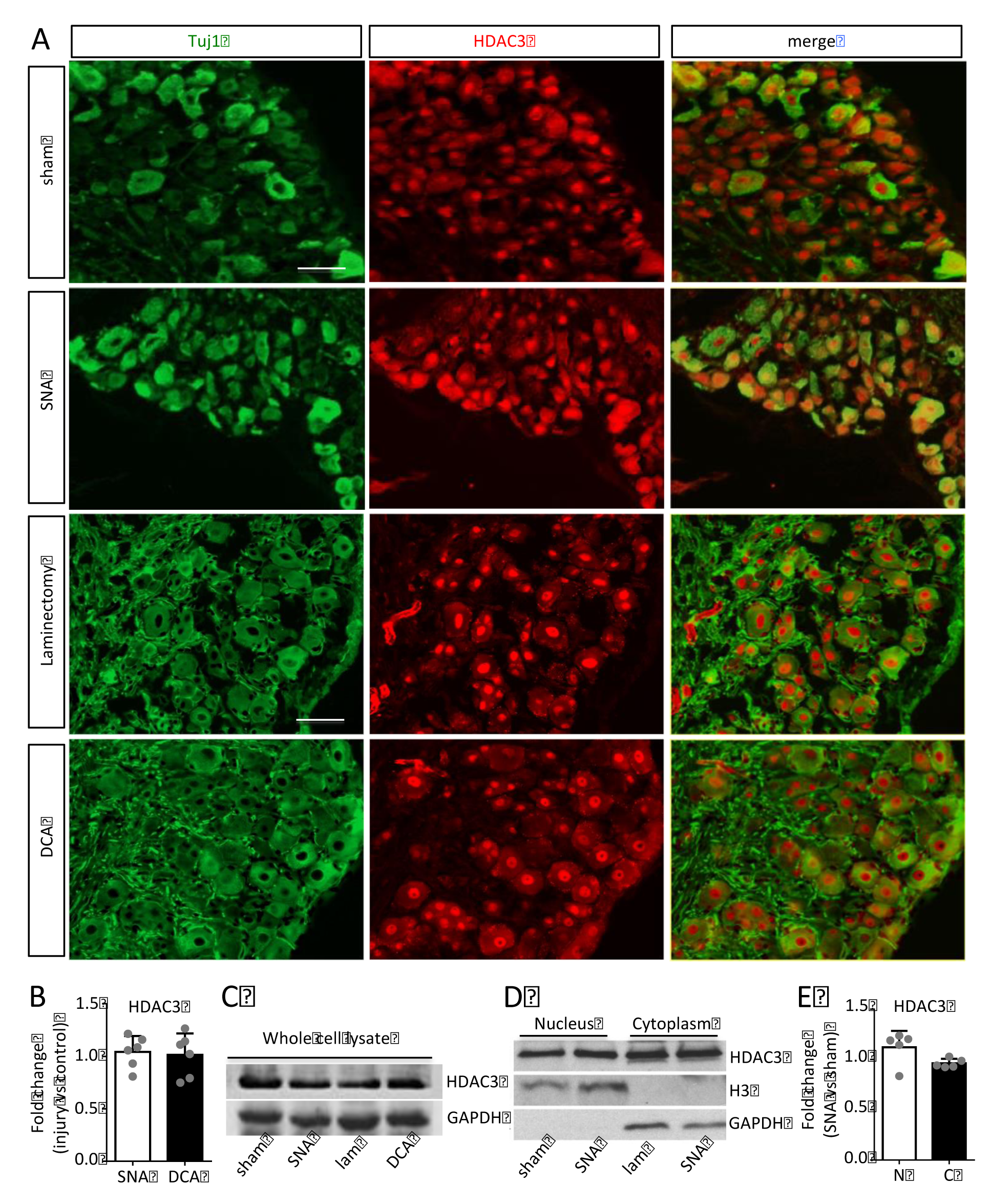
HDAC3 expression and subcellular localization in DRGs do not change after peripheral or central axonal injuries. A, B. Immunohistochemistry analysis of HDAC3 expression revealed that HDAC3 is highly expressed in lumbar DRG neurons (TUJ1 positive) and it does not change after SNA vs sham or DCA vs lam as shown by the bar graph after measurement of the HDAC3 immunofluorescent signal in TUJ1 positive neurons (B), N=4. Scale bar, (A), 100 μm. HDAC3 expression is mainly nuclear in DRG neurons, and its subcellular localization does not change after SNA. C. Immunoblot analysisshows that HDAC3 expression does not change after SNA or DCA. Data is expressed as mean fold change (band intensity) vs sham control ± s.e.m. N= 3 (Student’s t-test). D-E Immunoblot analysis of isolated nuclear and cytoplasmic DRG fractions shows that HDAC3 subcellular localization does not change after SNA. Data is expressed as mean fold change (band intensity) vs sham± s.e.m. N= 3 (Student’s t-test).

**Supplementary Figure 4.**
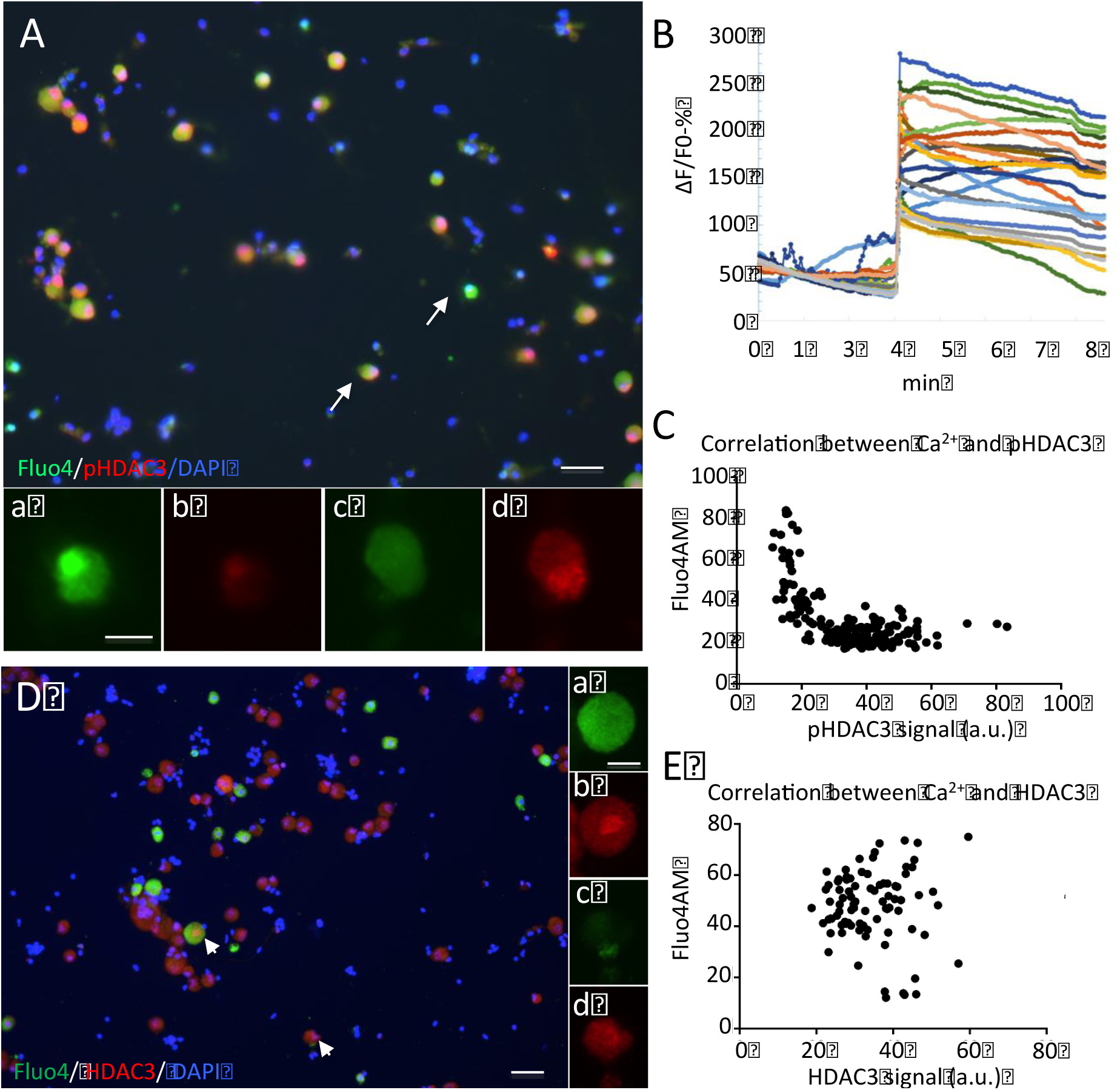
Dephosphorylation but not expression of HDAC3 is calcium dependent. A. pHDAC3 immunostaining in primary DRG neurons after KCl stimulation shows HDAC3 dephosphorylation specifically in DRG cells where high levels of calcium (Fluo4AM) were detected (arrows)(compare a-b with c-d). Scale bars, 300 μm (A) and 10 μm (a-d). B. Graphs show bursts of intracellular calcium during live imaging of Fluo4AM fluorescence after KCl administration. Data is expressed as single cell percentage of fluorescence variation versus basal fluorescence. C. Significant negative Pearson correlation between calcium levels (Fluo4AM) and pHDAC3 (r^2^: 0.4022); data is expressed as single cell fluorescence levels, N =176 cells from 3 biological replicates. D. HDAC3 immunostaining in primary DRG neurons after KCl stimulation shows similar HDAC3 levels in DRG cells where high and low levels of calcium (Fluo4AM) were detected (arrows). Scale bar, 300 μm and 10 μm (a-d). E. HDAC3 and Fluo4AM intensities show no correlation (r^2^: 0.00014); data is expressed as single cell fluorescence levels, N =86 cells from 3 biological replicates.

**Supplementary Figure 5.**
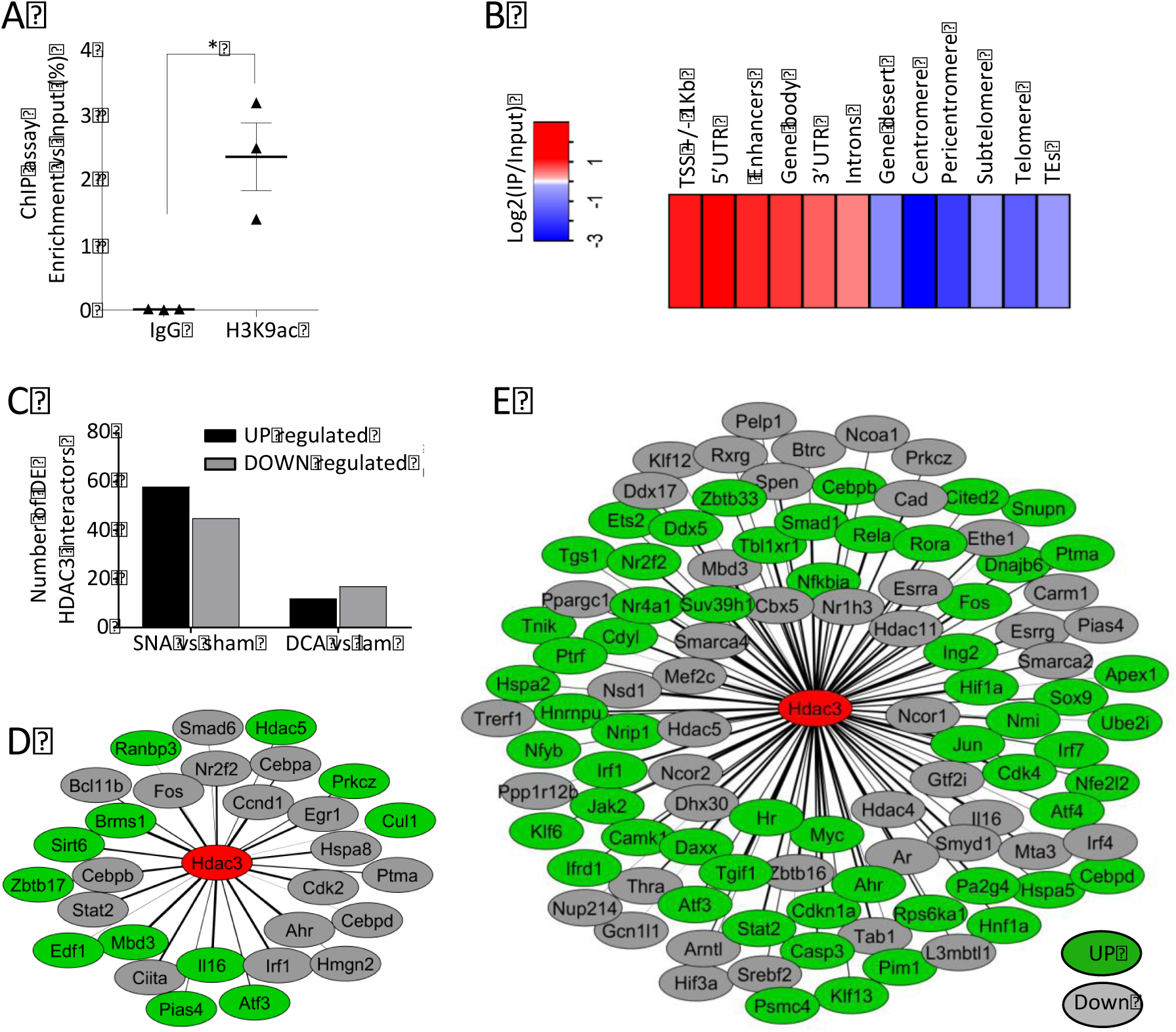
H3K9ac ChIPseq and HDAC3 interactome. A. Assessment of the enrichment of recovered DNA after H3K9ac ChIP versus control IgG ChIP. Data is expressed as percentage with respect to Input (average ± s.e.m. of 3 independent experiments, paired Student t-test, *p<0.05). B. Heatmap showing genomic distribution of H3K9ac reads, as Log_2_(IP vs Input) in a representative control sample. Note preferentially enrichment at TSS, enhancers and gene bodies. C. Bar graph shows the number of HDAC3 interactors identified using FpClass (score≥0.4) that are differentially expressed under the indicated conditions. Note that after SNA a higher percentage of HDAC3 interactors is differentially regulated with respect to DCA. D, E. Cytoscape visualization of the HDAC3 interactors that are differentially expressed upon DCA (D) or SNA (E). Upregulated interactors are in green, downregulated are in grey, HDAC3 is in red at the center of the network. The line thickness is proportional to the interaction score.

**Supplementary Figure 6.**
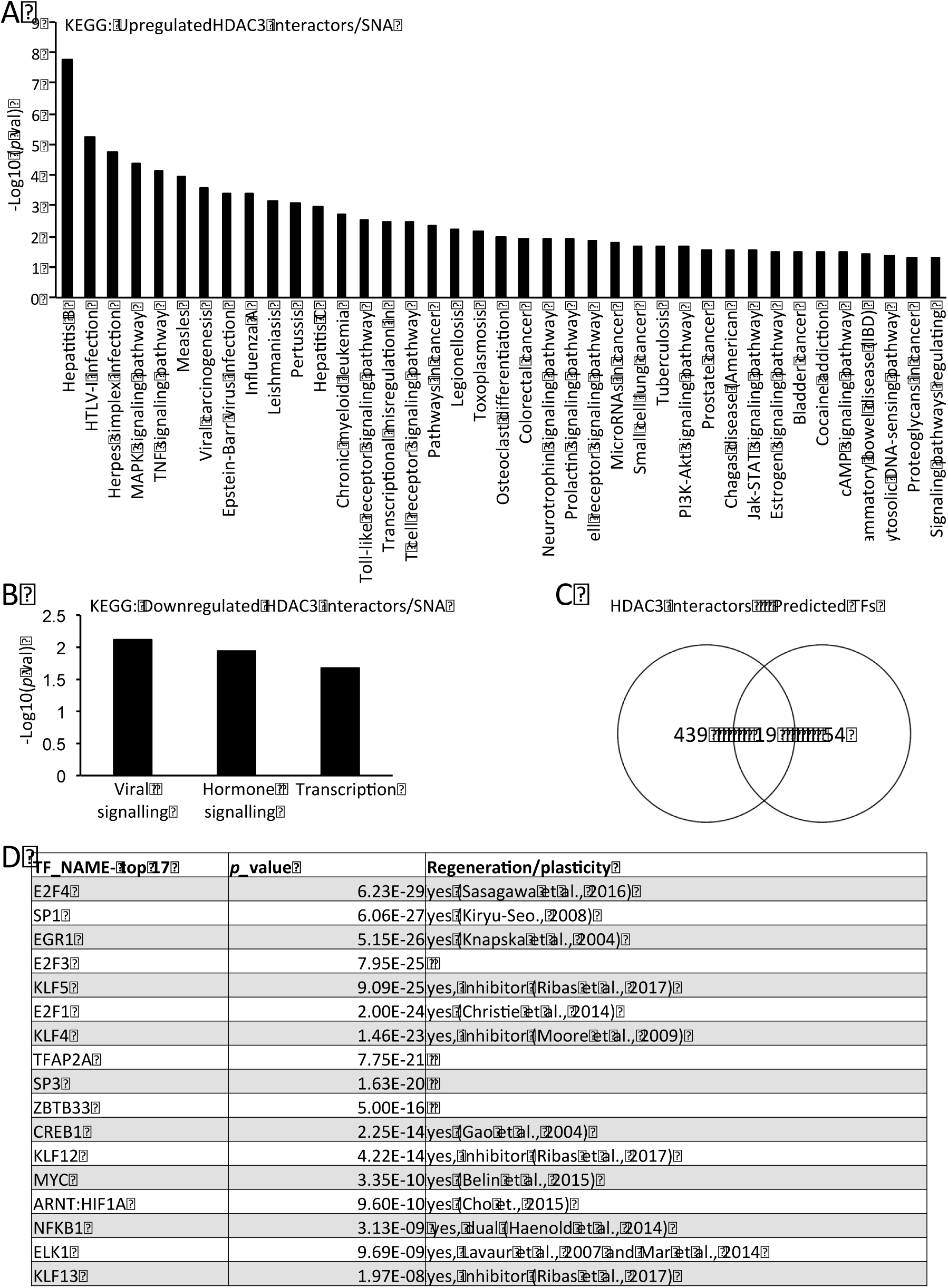
Upregulated HDAC3 interactors are related to regenerative signalling pathways. A. and B. Bar graph shows the KEGG pathway categories of the HDAC3 interactors sorted by −log_10_(*p* value) (cut off applied: *p*<0.05) that are up (A) or down (B) regulated after SNA. C. Venn diagram shows common transcription factors (TF) predicted to interact with HDAC3 (HDAC3 interactors) identified after *in silico* motif enrichment analysis of upregulated genes associated with increased H3K9ac after SNA following RNAseq (75 TF total) (see supplementary methods for details). D. Table shows a list of the identified TF predicted to interact with HDAC3, sorted by *p* value, and their previous involvement in regeneration/plasticity after axonal injury.

**Supplementary Figure 7.**
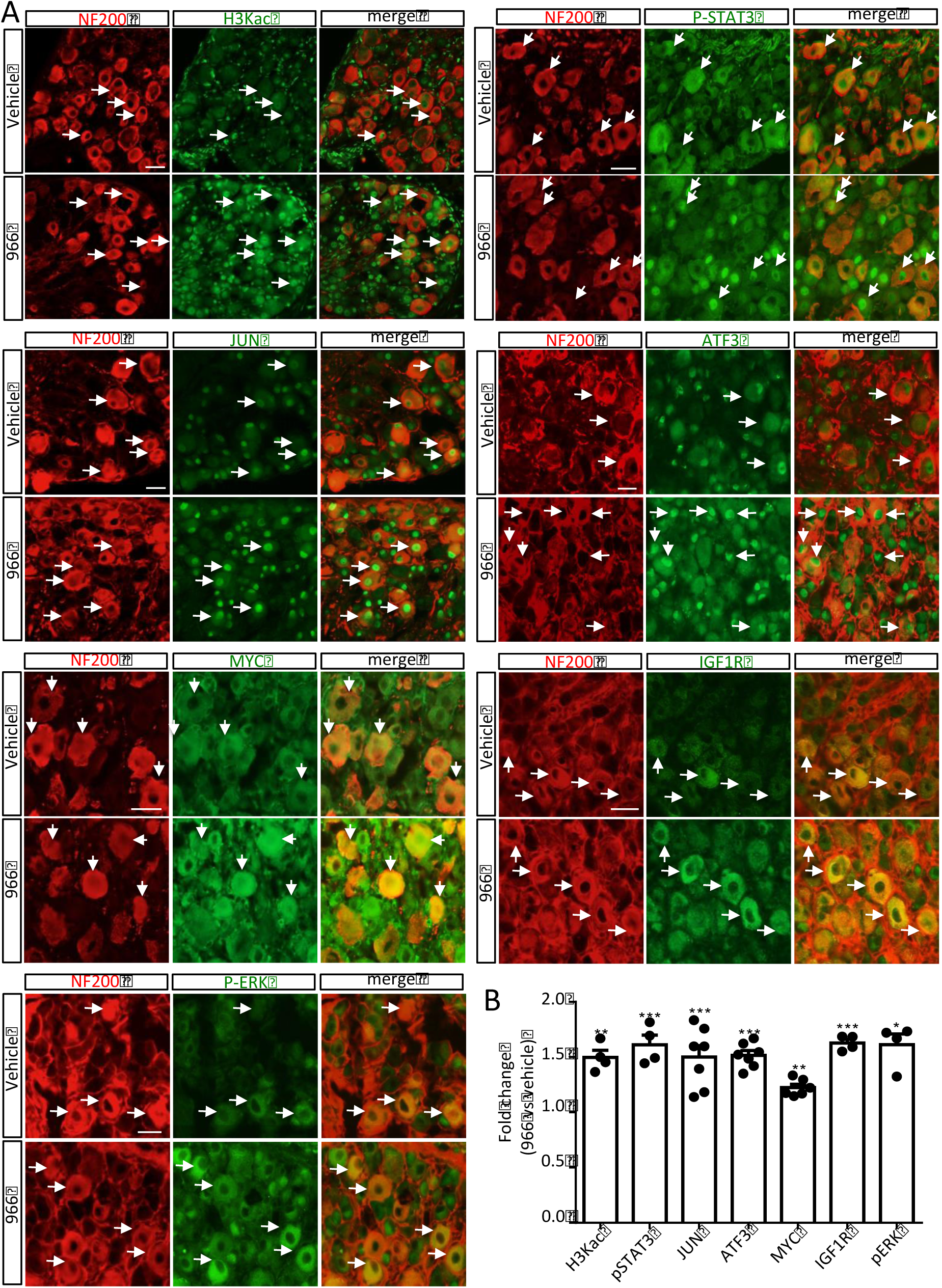
Intrathecal pharmacological inhibition of HDAC3 activity promotes H3K9ac and RAG expression in DRG neurons after spinal cord injury. A, B. Co-immunofluorescence of anti-neuronal and anti-regeneration associated proteins in DRG following SCI and treatment with 966 or vehicle. Shown is increased protein expression for regeneration-associated targets in NF200 positive DRG neurons (arrows) after 966 treatment (5 weeks following SCI). Scale bar, 50 μm. B. Data is expressed as fold change of fluorescence intensity in NF200^+^ cells, 966 versus vehicle ± s.e.m. N=5-7 * p<0.05, ** p<0.01, *** p<0.005 indicate significant difference versus vehicle (Student’s t-test).

**Supplementary Figure 8.**
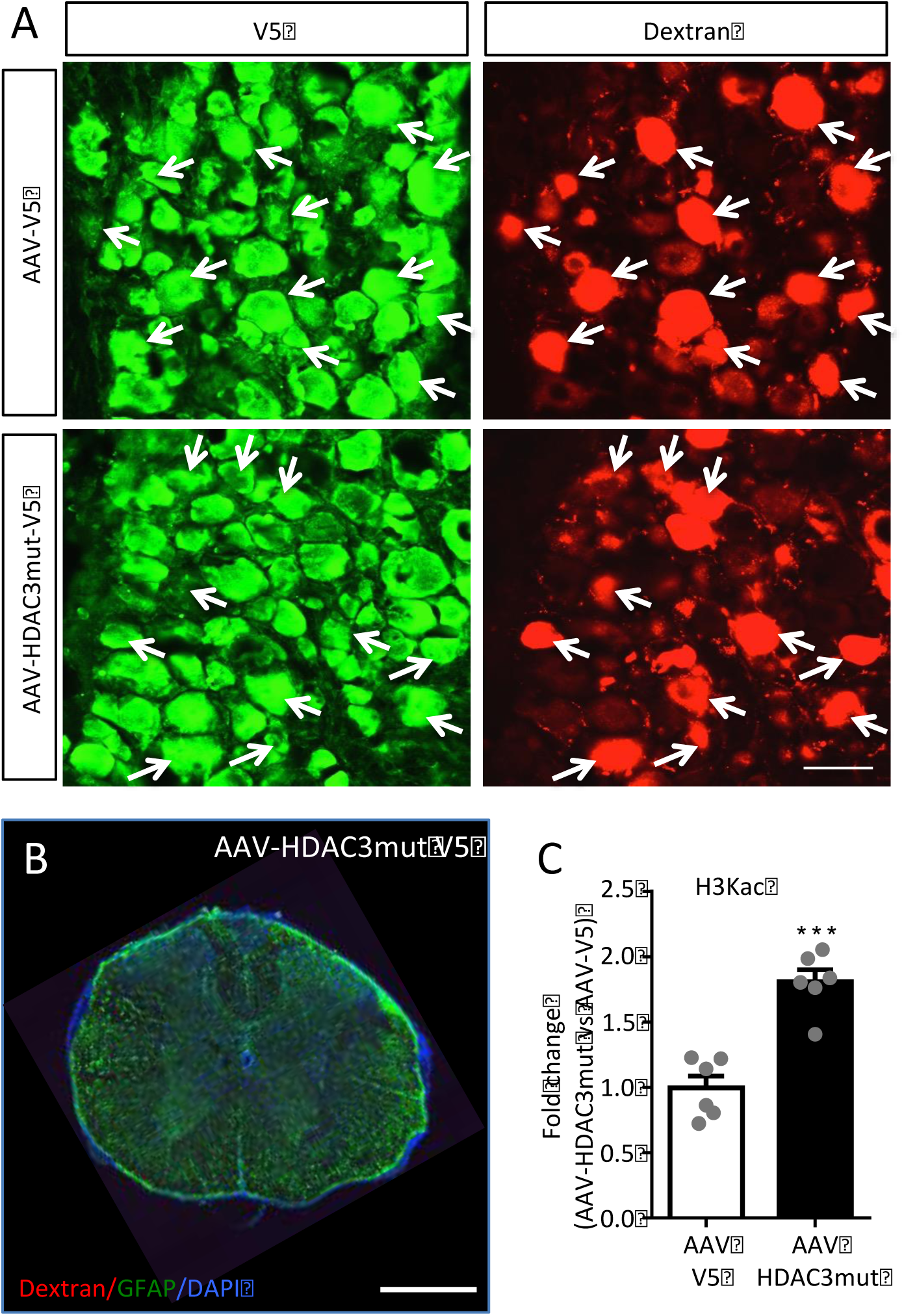
Genetic inactivation of HDAC3 deacetylase activity increases H3K9ac in DRGs. A. Immunofluorescence anti-V5 and dextran-red staining in whole DRG after sciatic nerve injection of AAV and dextran. Scale bar, 50μm. Arrows show co-localisation of V5-positive neurons in the majority of dextran-positive cells. B. Immunofluorescence anti-GFAP with fluorescent dextran and DAPI signal shows spinal cord coronal section of AAV-HDAC3 dead mutant injected mice 10 mm rostral from the lesion core, showing the absence of spared dextran+ axons. Scale bar, 500 μm C. Bar graph shows fold change of immunofluorescence intensity of H3K9ac in DRG after AAV-HDAC3mut vs control AAV± s.e.m. N= 6 animals per condition. (***p<0.005) indicate significant difference versus control AAV (Student’s t-test).

**Supplementary Figure 9.**
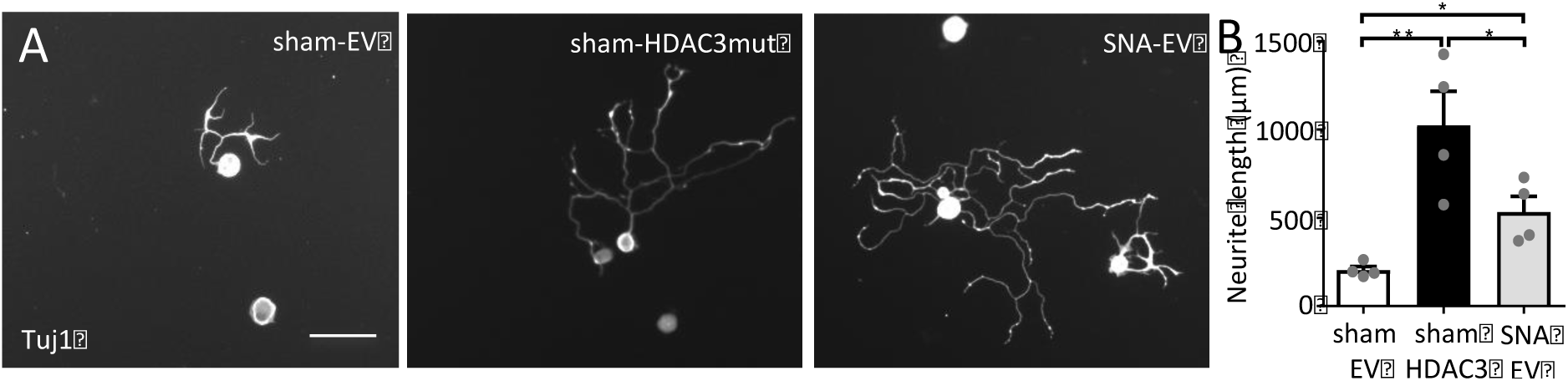
Intrathecal delivery of AAV-HDAC3-dead mutant induces neurite outgrowth of DRG neurons. A. AAV-mediated dead mutant HDAC3 *in vivo* infection of DRG neurons induced neurite outgrowth, including above the level of a conditioning lesion. Scale bars, 100 μm. B. Data is expressed as mean neurite length ± s.e.m. n= 4 technical replicates from 3 biological replicates. (*p<0.05, **p<0.01) indicate significant difference versus sham+EV (Empty Vector) (ANOVA followed by Bonferroni test).

**Supplementary Figure 10.**
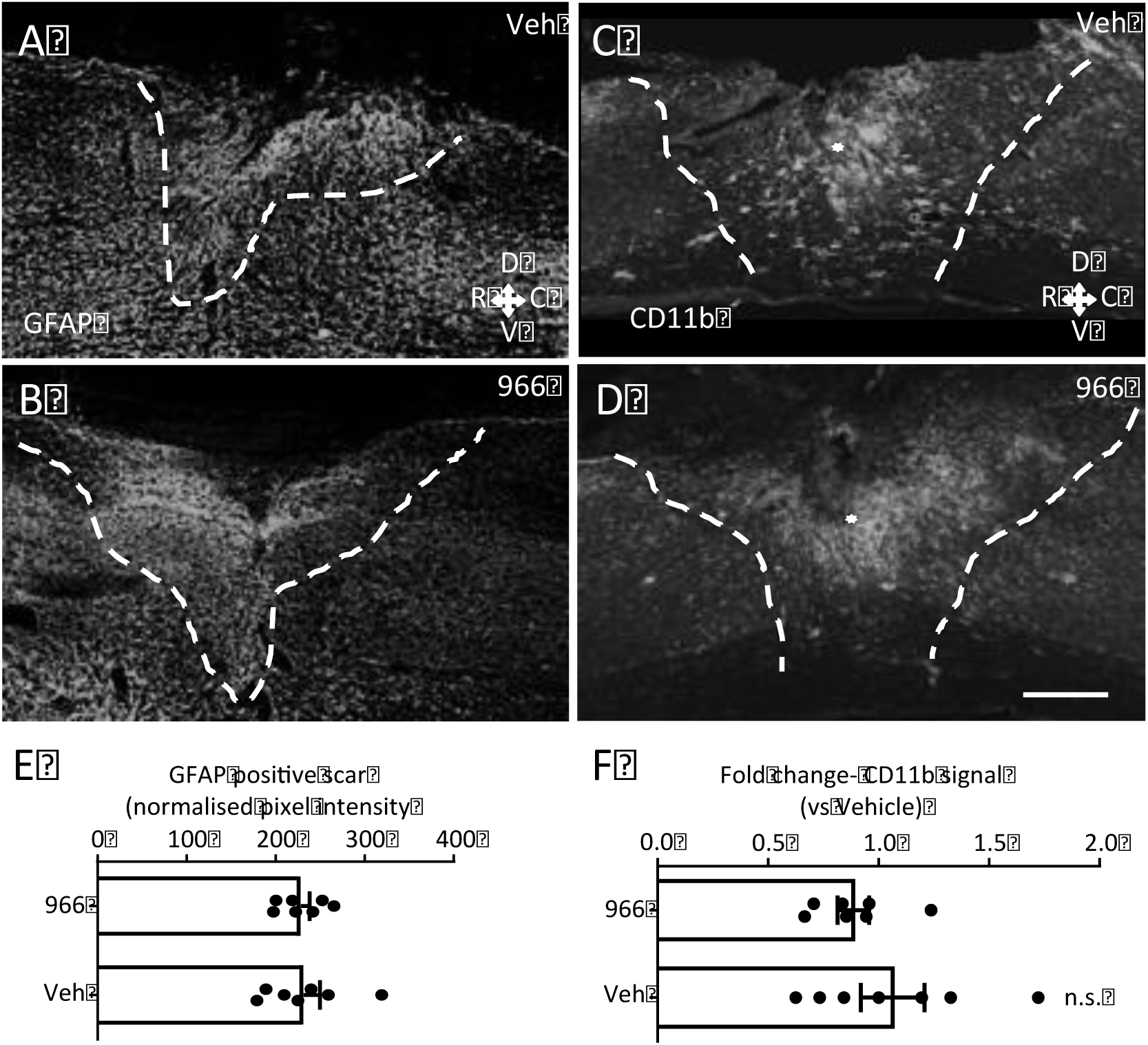
GFAP and CD11b positive immunoreactivity around the injury site. A-D. Immunofluorescence anti-GFAP (A, B) or anti-CD11b (C, D) shows that the GFAP positive scar (five weeks after SCI) (A, B) or CD11b immunoreactivity (C, D) around the injury site are not affected by intrathecal administration of RGFP966 (966) through osmotic minipump vs vehicle. Scale bar, 500 μm. E. Data is expressed as fold change of GFAP^+^ area pixel intensity (dotted lines define boundaries of GFAP^+^ scar) between 966 vs vehicle ± s.e.m. N= 7. (Student’s t-test). F. Data is expressed as fold change of CD11b^+^ pixel intensity (within the glial scar area, dotted lines) between 966 vs vehicle ± s.e.m. N= 7 (veh) and 9 (966). (Student’s t-test).

**Supplementary Figure 11.**
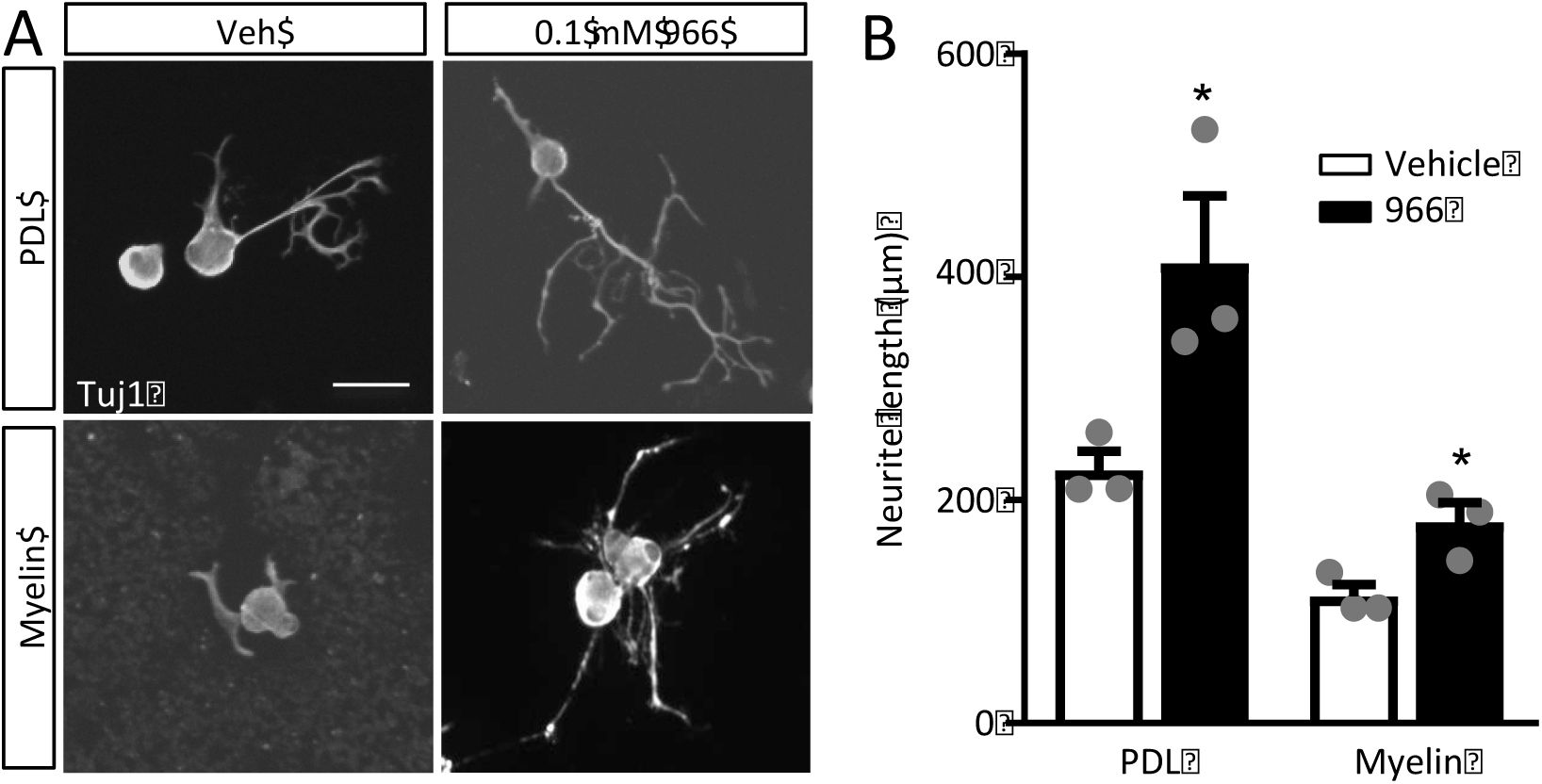
Intrathecal pharmacologic inhibition of HDAC3 activity induces neurite outgrowth of DRG neurons on permissive and inhibitory substrates. A. Intrathecal administration of RGFP966 (966) *in vivo* induced neurite outgrowth on PDL and myelin substrates. Scale bar, 100 μm. B. Data is expressed as mean neurite length ± s.e.m. N=3 technical replicates from 3 biological replicates (animals). (\**p*<0.05) indicate significant difference versus respective veh (Student’s t-test).

**Suppl File 1**: Complete dataset of RNAseq, and ChIPseq read intensities in each indicated condition at the level of TSS+/-1000bp and gene body. For each gene, the gene Ensembl ID, the gene name, the chromosome coordinates, the logFC and *p* value of gene expression and H3K9ac occupancy are reported.

**Suppl. File 2**: List of HDAC3 interactors identified with FpClass (score >0.4) and expressed in the DRG RNAseq dataset.

**Suppl. File 3**: GO and KEGG analysis of the HDAC3 interactors Up and Down-regulated after SNA or DCA. In yellow the significant terms (*p* value <0.05)

**Suppl. File 4**: Transcription factor enrichment analysis on the H3K9ac dependent Upregulated genes after SNA, using Pscan (Jaspar 2016, TSS −450/+50bp). In yellow the statistically significant TFs (Bonferroni *p* value <0.05).

**Suppl. File 5**: GO and KEGG analysis of the HDAC3 network. In yellow the significant terms (*p* value <0.05)

